# Cortical excitability inversely modulates fMRI connectivity via low-frequency neuronal coupling

**DOI:** 10.64898/2026.03.12.710517

**Authors:** David Sastre-Yagüe, Simone Blanco Malerba, Federico Rocchi, Silvia Gini, Gabriele Mancini, Alexia Stuefer, Ludovico Coletta, Shahryar Noei, Marija Markicevic, Filomena Grazia Alvino, Valerio Zerbi, Alberto Galbusera, Jean Charles Mariani, Stefano Panzeri, Alessandro Gozzi

**Affiliations:** Functional Neuroimaging Laboratory, Center for Neuroscience and Cognitive systems, Istituto Italiano di Tecnologia, Rovereto, Italy; Center for Mind and Brain Sciences, University of Trento, Rovereto, Italy; Institute for Neural Information Processing, Center for Molecular Neurobiology (ZMNH), University Medical Center Hamburg-Eppendorf (UKE), Hamburg, Germany; Optical Approaches to Brain Function Laboratory, Istituto Italiano di Tecnologia (IIT), Italy; Department of Pharmacy and Biotechnology, University of Bologna, Bologna, Italy; Neuroinformatics Laboratory (NiLab), Bruno Kessler Foundation (FBK), Trento, Italy; Department of Neurology, Inselspital, University Hospital and University of Bern, 3010 Bern, Switzerland; Department of Psychiatry, Faculty of Medicine, University of Geneva, Switzerland; Department of Basic Neurosciences, Faculty of Medicine, University of Geneva, Switzerland

## Abstract

The neural mechanisms supporting fMRI connectivity are poorly understood. Leveraging an aggregated analysis of novel and existing chemogenetic manipulations and electrophysiological recordings in the mouse medial prefrontal cortex (PFC), we show that local cortical excitability inversely modulates large-scale fMRI connectivity. Specifically, we find that bidirectional chemogenetic manipulations of cortical excitability produce opposite effects on fMRI connectivity, resulting in fMRI hypoconnectivity when excitability is enhanced, and fMRI hyperconnectivity when excitability is suppressed, despite corresponding increases or decreases in local neuronal firing. Notably, while each chemogenetic manipulation produces a distinct profile of interareal electrophysiological coherence, only low-frequency (< 4Hz) coherence predicts the direction and magnitude of the ensuing fMRI connectivity changes. Biophysical network modelling shows that the observed electrophysiological coherence profiles can arise from the interaction between direct inter-area communication changes mediated by local firing-rate variations, and shared larger-scale low-frequency covariation of ongoing neuronal activity. Together, our results reveal an inverse relationship between regional cortical excitability and large-scale fMRI connectivity, and indicate that fMRI connectivity is primarily supported by low frequency (<4 Hz) electrophysiological coupling. These findings open new avenues for modeling and interpreting fMRI connectivity in health, and in response to pathological or exogenous perturbations linked to cortical excitability.

## Introduction

Spontaneous brain activity mapping via fMRI provides a reliable, non-invasive window into the macroscopic organization of the human brain^1^. Highly structured spatiotemporal patterns of fMRI activity, known as resting-state networks (RSN), can be reliably identified across individuals, providing a basis for probing how large-scale networks evolve across development^2^, support cognition^3, 4^, adapt through learning^5^, and are altered in neuropathology (Menon, 2011; Pagani et al., 2021). Within this framework, synchronous fMRI activity within and across RSNs, termed ‘functional connectivity’, is often interpreted as an index of large-scale neuronal communication between brain regions.

While stimulus- or task-evoked fMRI activity is strongly associated with local increases in γ-band and higher-frequency neuronal activity^6, 7, 8, 9, 10^, the neural mechanisms supporting interareal resting-state fMRI connectivity remain a matter of active debate. Previous studies have associated fMRI connectivity with a broader range of timescales and frequency bands, including band-limited γ activity ^9, 10, 11^, δ oscillations^12, 13, 14^ and slow (< 1 Hz)^13, 15, 16^ or ultra-slow (<0.1 Hz) fluctuations^17, 18, 19, 20^. As a result, how the interplay between ongoing neural rhythms and local neuronal activity supports and influences fMRI connectivity remains an open question.

Perturbational studies provide a powerful route to causally disentangle the relationship between local and global-scale fMRI dynamics, and its underlying neural activity^21, 22, 23^. Within this framework, chemogenetic approaches (chemo-fMRI^24^) offer the opportunity to durably modulate brain activity, thereby enabling robust estimation of fMRI connectivity under steady-state post-manipulation conditions^25^. However, chemo-fMRI studies have only rarely been anchored to direct measurements of the underlying neuronal population activity^16, 23^. This gap is particularly critical in light of recent work showing that the net *in vivo* effects of chemogenetic perturbations cannot be reliably predicted by receptor design alone, but can even oppose the intended direction of modulation, depending on expression levels, targeted cell types, promoter used for viral expression, and duration of activation^26, 27, 28^. These observations raise the possibility that the heterogeneous effect of chemogenetic manipulations on fMRI connectivity can be reconciled along a single dominant neurophysiological axis once their effects on neural population activity are experimentally assessed using *in vivo* electrophysiological recordings.

In the present work, we posit that cortical excitability, an operational readout of local E/I balance, provides a dominant neurophysiological axis that shapes interareal fMRI connectivity and reconciles how distinct oscillatory coupling profiles map onto common connectivity outcomes. In recurrent cortical circuits, excitation and inhibition are tightly coupled, and multiple microcircuit parameters (including excitatory and inhibitory synaptic strength, and the intrinsic excitability of each population) jointly determine the circuit’s operating point (i.e., its circuit-level excitability^29^). Theorethical work suggests that this operating point can be indexed *vivo* by the level of average population firing rate at which the local circuit operates^29, 30^. We therefore hypothesize that shifts in this circuit-level excitability, whether induced by perturbations, or arising spontaneously across development, brain states, and pathology, can systematically modulate interareal synchronization and, in turn, fMRI connectivity. This framework is consistent with the established role of E/I balance in both setting the dominant frequencies of local oscillations and in shaping local and long-range neuronal synchronization^31, 32, 33, 34^. It is also consistent with evidence that altered local E/I balance frequently co-occurs with disrupted fMRI connectivity in animal models as well as in human developmental disorders^30, 35, 36, 37^. Importantly, this hypothesis is directly testable using chemogenetic perturbations combined with *in vivo* recordings of local spiking activity. Within this framework, DREADD-based chemogenetics^25^ enables remote, receptor-specific modulation via G-protein-coupled signaling which can influence both intrinsic excitability and synaptic transmission ^38^. Leveraging these properties, chemogenetic targeting of distinct circuit components provides a principled way to drive different firing-rate regimes and test how such shifts reshape fMRI connectivity.

Here, we use chemo-fMRI and multielectrode electrophysiology in the mouse medial prefrontal cortex (PFC) to test the hypothesis that circuit-level excitability acts as systems-level physiological axis that controls fMRI connectivity, and accounts for its relationship with spontaneous electrophysiological oscillations. By aggregating novel and previously published cell-type specific chemogenetic manipulations into a unified analytical framework, we examine how distinct local perturbations of circuit-level excitability reshape fMRI connectivity and interareal electrophysiological coupling at different frequencies. Across perturbations, we find that large-scale fMRI connectivity varies inversely with local cortical excitability, and tracks with low-frequency (<4 Hz) interareal electrophysiological coupling. These findings provide a parsimonious mechanistic framework for interpreting and modeling fMRI connectivity in the context of perturbations linked to cortical excitability.

## Results

### Chemogenetic activation of pyramidal neurons suppresses cortical rhythmic activity

To causally test whether and how DREADD-induced changes in neocortical excitability affect large-scale fMRI connectivity, we first increased the intrinsic excitability of pyramidal neurons in the mouse PFC by virally transducing the hM3Dq receptor under the expression of the CamkII-α promoter. We hereafter refer to this manipulation as ↑Excitation. A schematic of our experimental design is shown in Figure 1a.

**Figure 1.**
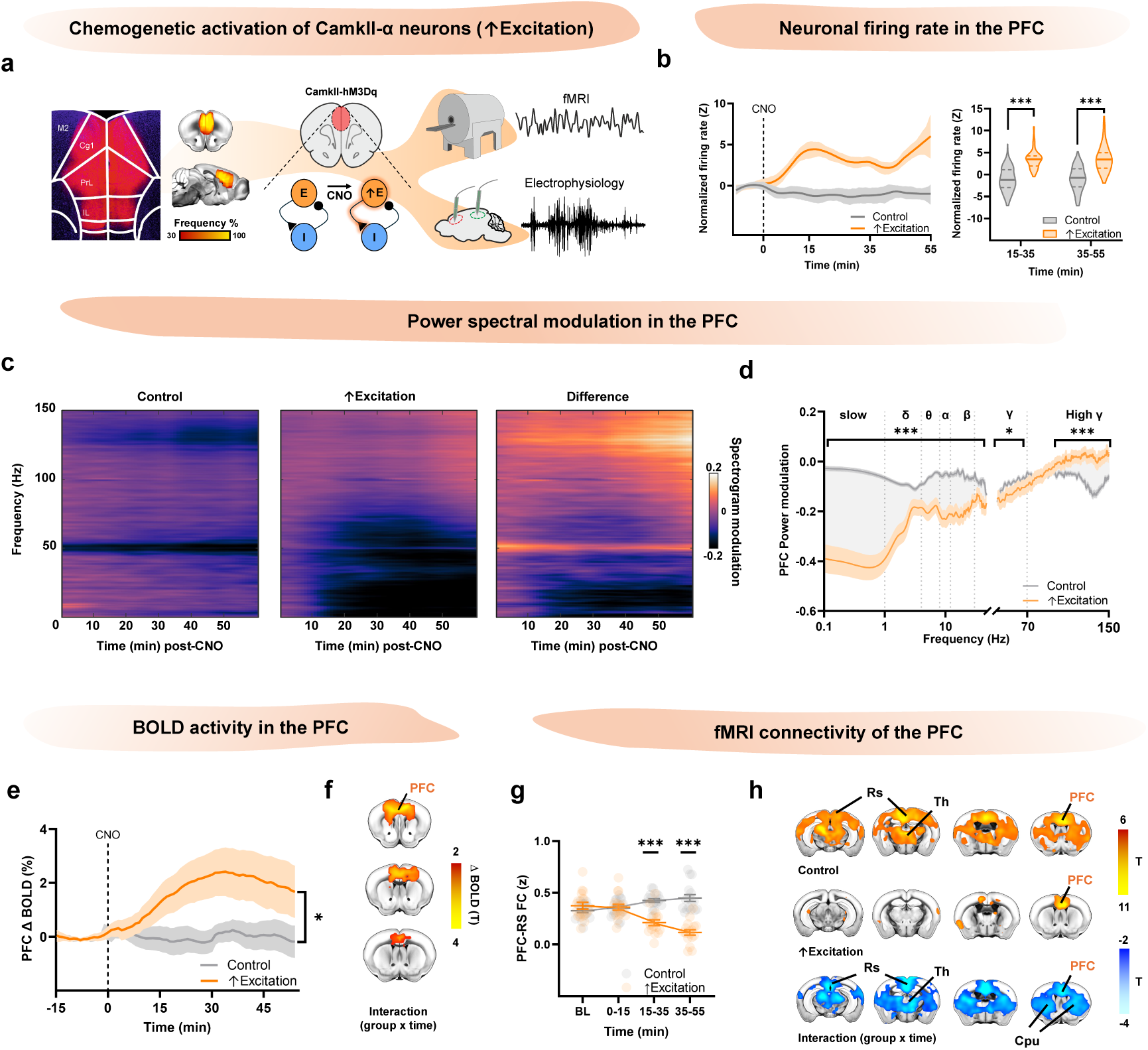
Chemogenetic activation of pyramidal neurons (↑Excitation) increases local excitability and induces fMRI hypoconnectivity. **a)** Experimental design. CamkIIα-hM3Dq receptor was injected bilaterally in the PFC. Leftmost panels show representative viral expression and group-level viral coverage maps across animals. The effect of chemogenetic activation was mapped using fMRI and multielectrode electrophysiology. **b)** Baseline-normalized firing rate in the PFC (right, summary over 15-35 and 35-55 min; Wilcoxon rank-sum test; ↑Excitation: 100 non-overlapping time bins from n=5 mice; control: 140 bins from n=7 mice). **c)** Group-average spectrogram modulation in PFC after CNO administration (control n=7; ↑Excitation n=6), and difference (↑Excitation-control, right). **d)** Frequency resolved spectral modulation after chemogenetic activation of pyramidal neurons (n=120 and n=140 bins from n=6 ↑Excitation and n=7 control mice, Student’s t-tests, cluster FWER corrected). **e)** Average fMRI BOLD signal change in PFC after CNO administration (↑Excitation and control, n=17 each, group x time interaction). **f)** Voxel-wise mapping of BOLD signal change elicited by ↑Excitation over the maximal effect window (20-35 minutes post CNO, group x time interaction, |t|>2.1, cluster FWER corrected). **g)** fMRI connectivity between PFC and Rs across time windows (↑Excitation and control, n= 17 mice each, group x time interaction, linear mixed-effects model). **h)** fMRI connectivity of the PFC in the window of maximal connectivity difference (35 to 55 minutes post CNO, group x time interaction, cluster corrected, ↑Excitation n=17, control n=17). BL: baseline; PFC, prefrontal cortex; Rs, retrosplenial cortex; Th, thalamus; CPu, caudate-putamen. Normalized firing rate timeseries are shown as mean ± C.I. of loess fit. All other line plots show mean ± SEM; *p<0.05, ***p<0.001. Violin plots show median (thick line) and 75th and 25th percentiles (dashed line).

We first validated the effectiveness of this manipulation by recording electrophysiological activity in the PFC before and after peripheral administration of the DREADD actuator clozapine-N-oxide (CNO) (Figure 1a). In this and subsequent DREADD experiments, we recorded both local field potentials (LFPs) and multi-unit activity (MUA). LFPs were used to quantify local oscillatory activity and frequency-resolved interareal coupling, which we subsequently related to changes in fMRI connectivity. MUA provides instead a population-level measure of average local firing, i.e., our operational readout of circuit-level excitability in the manipulated region. This choice is supported by theoretical and experimental work showing that population firing captures the net effect of multiple factors governing E/I interactions, including E-I synaptic strengths as well as the intrinsic excitability of each population. Because excitation and inhibition are tightly coupled in cortex, manipulations that increase (or decrease) excitatory drive often elevate (or decrease) firing in both excitatory and inhibitory populations. Consistent with this, *in silico* simulations (Figure S1B) show that increasing (or decreasing) the intrinsic excitability of excitatory neurons increases (or decreases) both excitatory and inhibitory-neuron firing. Similarly, strengthening excitatory synapses elevates firing in both excitatory and inhibitory populations (Figure S1C). Average population firing thus captures the net effect of E/I interactions, and indexes the excitability level at which the circuit operates^30^. Under this operationalization, in this work DREADD-induced increases (or decreases) in cortical firing as measured by MUA reflect heightened (or reduced) cortical excitability.

Consistent with this framework, chemogenetic activation of CamkIIα^+^ neurons led to a robust and sustained increase in firing activity in the manipulated area (Figure 1b, p<0.001, Wilcoxon rank-sum test, FDR corrected), indicating heightened cortical excitability. Local field potential (LFP) spectrograms obtained from the same electrophysiological recordings revealed that this increase in neuronal firing was accompanied by a marked suppression of rhythmic activity extending up to the γ range (<70 Hz, Figure 1c,d, cluster1=0.1-40 Hz, p<0.001, cluster2=52-70 Hz, p=0.047, cluster-based permutation test), and a concomitant increase in high-γ power (70-150 Hz, Figure 1c,d, 90-150 Hz, p < 0.001, cluster-based permutation test). The latter finding is consistent with the observed MUA increase, since high-γ power, also referred to as broadband high frequency activity, is an established correlate of local neuronal firing^39^. Taken together, these results show that chemogenetic activation of cortical pyramidal neurons increases local circuit-level excitability (as reflected by elevated firing rate), while strongly suppressing rhythmic activity in the manipulated area.

Previous research has shown that optogenetically increasing cortical excitation-inhibition (E/I) ratio in the mouse PFC impairs social function^40^. To test whether this was also the case for our chemogenetic manipulation, we conducted a three-chamber sociability test in control and hM3Dq-expressing mice. Consistent with previous findings, we found that ↑Excitation led to a robust reduction of mouse sociability in manipulated mice (Figure S2b, sociability index, p=0.009, Student’s t-test). Together, these results suggest that the employed chemogenetic manipulation is both physiologically effective and behaviorally relevant.

### Chemogenetic activation of pyramidal neurons results in fMRI hypoconnectivity

To investigate the impact of ↑Excitation on large-scale fMRI connectivity, we conducted resting-state fMRI recordings in control and hM3Dq-expressing mice following administration of DREADD-activator CNO (Figure 1). We first investigated whether ↑Excitation modulates local BOLD activity (Figure 1e-f). Consistent with the established association between high-frequency LFP power and local BOLD fMRI activity^8, 9, 10^, and the results of our electrophysiological recordings in hM3Dq-expressing mice, we found a significant increase in BOLD signal in the PFC of manipulated mice. The effect reached its peak approximately 30 minutes after CNO administration (Figure 1e, p=0.041, group x time interaction, Figure 1f, group x time interaction, |t|>2.1, cluster-corrected). Conversely, no significant BOLD signal changes were detected in control mice receiving CNO. These findings suggest that increased neuronal firing, reflecting heightened cortical excitability, is accompanied by local increases in BOLD activity.

We next asked how increased cortical excitability affects fMRI connectivity. To address this question, we conducted a seed-based analysis to probe fMRI connectivity of the PFC in control and hM3Dq-expressing animals following CNO administration (Figure 1g-h). We found that ↑Excitation robustly reduced fMRI connectivity between the manipulated region (PFC) and several of its targets within the default mode network (DMN), including the striatum, thalamus, and retrosplenial cortex (Figure 1h, group x time interaction, linear mixed-effects model, |t|>2.1, p<0.05, cluster-corrected). The effect reached its peak amplitude between 35 and 55 minutes post-CNO injection (Figure S3a and Figure 1g, p<0.001, group x time interaction, FWER corrected). These results show that increasing local circuit-level excitability counterintuitively decreases fMRI connectivity of the affected cortical region, despite locally increased neuronal firing and BOLD activity.

We next performed a set of control experiments to probe the specificity and generalizability of this finding. We first tested whether the observed reduction in fMRI connectivity could be a indirect consequence of a local ceiling effect in the hyperemic response induced by ↑Excitation. In this scenario, any manipulation that produces a large (ceiling), sustained hyperemic response in PFC, irrespective of its origin, should suppress intrinsic BOLD signal fluctuations and thereby reduce functional connectivity of this region. To test this hypothesis, we analyzed fMRI connectivity in mice receiving the NMDAR antagonist phencyclidine (PCP; Figure S4). We did so because we previously found^41^ that PCP administration elicits a robust local BOLD response in the mouse PFC, approximately twice as large as that observed here (5% vs 2%, Figure S4a). If a hyperemia in the PFC alone were sufficient to suppress intrinsic fMRI signal fluctuations (and thus reduce connectivity) through purely vascular mechanisms, PCP-treated mice should exhibit connectivity disruptions comparable to (or greater than) those induced by ↑Excitation. This was not the case, as we found PFC connectivity to be broadly preserved in mice administered with PCP (Figure S4b, |t|>2.1, p<0.05 FWER-corrected). This result argues against a purely vascular origin for the fMRI hypoconnectivity induced by ↑Excitation.

We next asked whether ↑Excitation-induced hypoconnectivity, initially observed under light sedation, could also be detected under awake conditions. To this end, we carried out fMRI mapping in a second cohort of control and DREADD-expressing mice scanned under awake, head-fixed conditions^42^. We specifically examined whether CNO administration in ↑Excitation group would reduce fMRI connectivity between the PFC and its target regions (Figure S5a, p=0.003, group x time interaction, Figure S5b, group x time interaction, t>1.7, p<0.05, FWER-corrected). Although the magnitude of this effect was weaker than that observed under sedation, a finding consistent with a more desynchronized cortical state in aroused animals^42, 43^, hM3Dq-expressing mice exhibited reduced fMRI connectivity in the medial thalamus, a key node of the DMN. Additional foci of decreased connectivity that did not survive cluster-level significance were also present in other DMN components, such as the retrosplenial cortex (Figure S5b). To statistically corroborate these preliminary observations, we next performed an aggregate analysis combining sedation and awake datasets using a combined linear mixed-effects model. Group (↑Excitation vs. control), time window (baseline vs. post-CNO), and experimental condition (sedation vs. awake) were included as fixed effects, with subjects modeled as random factors. This approach allowed us to test for group × time interactions while accounting for variability produced by the different brain state across recordings (i.e., sedation vs. awake). Consistent with a shared direction of ↑Excitation-induced connectivity changes across brain states, this analysis revealed a robust group × time interaction in multiple DMN regions, including thalamus, retrosplenial cortex, striatum, with manipulated mice showing fMRI hypoconnectivity compared to controls in those areas (Figure S5c, p<0.001, S4d, t>2.1, p<0.05, cluster-corrected).

Importantly, existing chemo-fMRI work suggests that ↑Excitation-induced hypoconnectivity may not be specific to polymodal cortical regions like the PFC, but may also extend to unimodal sensory cortices. In a previous study, cortical excitability was increased via CamkIIα-driven hM3Dq expression in primary somatosensory cortex under the same sedation protocol used here, and reduced fMRI connectivity was reported in the manipulated regions in good agreement with our PFC results^44^. To enable direct comparability with the experiments reported here, we re-analyzed those fMRI data within the present analytical framework and observed robust fMRI hypoconnectivity both in contralateral and ipsilateral targets of the manipulated area (Figure S6a, group x time interaction, |t|>2.1, cluster -corrected). These convergent findings suggest that increasing cortical excitability can lead to fMRI hypoconnectivity both in unimodal and polymodal cortical regions.

### Chemogenetic inhibition of PV^+^ interneurons enhances cortical oscillatory activity

Our ↑Excitation experiments revealed an fMRI signature characterized by locally increased BOLD activity in the manipulated region, and reduced long-range fMRI connectivity between the manipulated region and its long-range projection targets. However, cortical excitability can also be increased through reduced inhibition. Does this manipulation similarly affect fMRI connectivity? To address this question, we employed DREADD-based chemogenetics to inhibit PV^+^ GABAergic interneurons in the mouse PFC. This was achieved by transducing a Cre-dependent version of the inhibitory DREADD receptor hM4Di in PV::cre mice (Figure 2a). We hereafter refer to this manipulation as ↓Inhibition.

**Figure 2.**
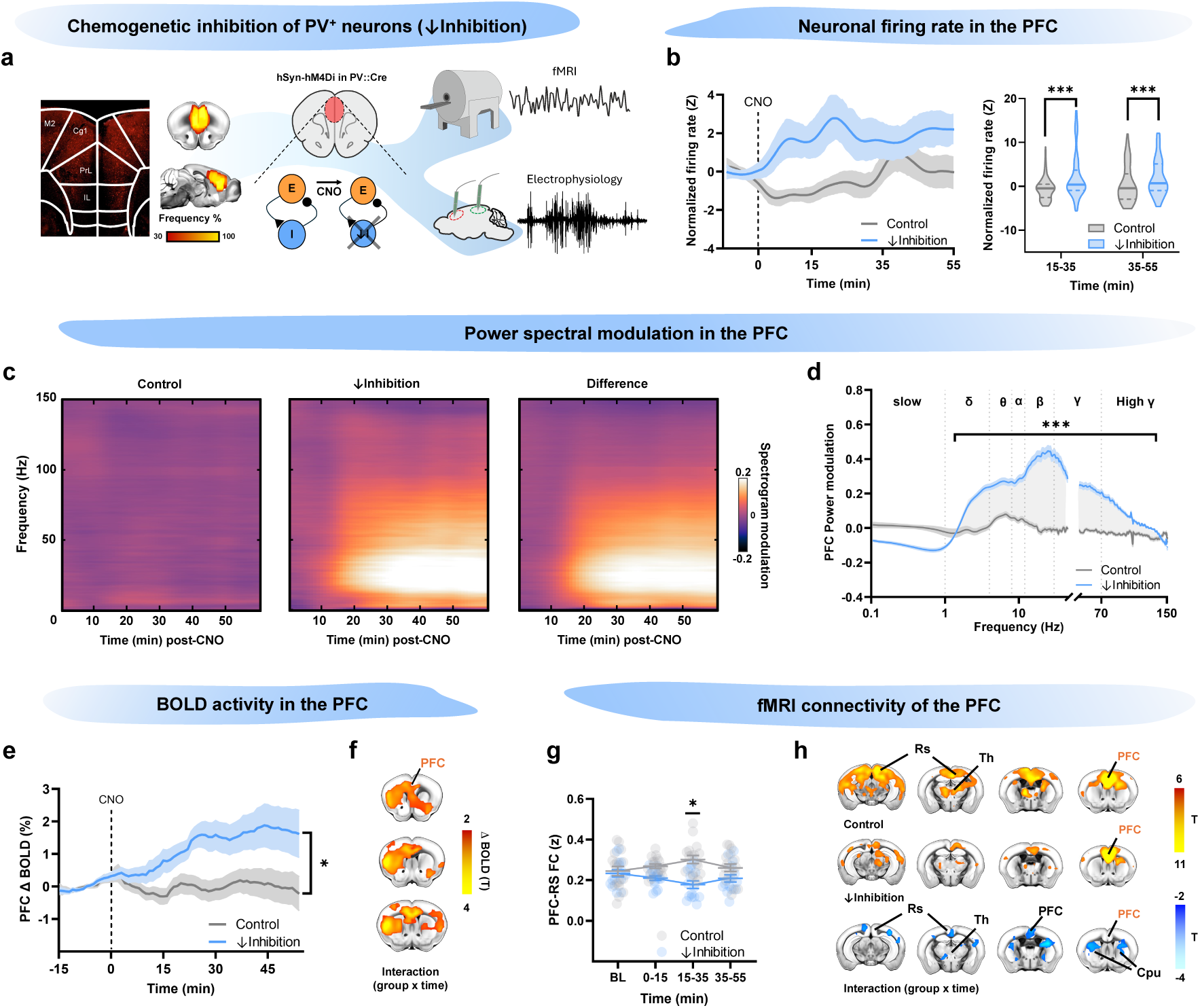
Chemogenetic inhibition of PV+ interneurons (↓Inhibition) increases local excitability and induces fMRI hypoconnectivity. **a)** Experimental design. hSyn-DIO-hM4Di-mCherry was injected bilaterally in the PFC of PV::Cre mice. Leftmost panels show representative viral expression and group-level viral coverage maps across animals. The effect of chemogenetic inhibition was mapped using fMRI and multielectrode electrophysiology. **b)** Baseline-normalized firing rate in the PFC (right, summary over 15–35 and 35-55 min; Wilcoxon rank-sum test; ↓Inhibition: 220 non-overlapping time bins from n=11 mice; control: 200 bins from n=10 mice). **c)** Group-average spectrogram modulation in PFC after CNO administration (control n=10; ↓Inhibition n=11), and corresponding difference (↓Inhibition - control, right). **d)** Frequency resolved spectral modulation after chemogenetic inhibition of PV+ interneurons (n=220 and n=200 bins from n=11 ↓Inhibition and n=10 control mice, respectively, Student’s t-tests, cluster FWER-corrected). **e)** Average fMRI BOLD signal change in PFC after CNO administration (↓Inhibition n=15; control n=17, group × time interaction). **f)** Voxel-wise mapping of BOLD signal change elicited by ↓Inhibition over the maximal effect window (20-35 minutes post CNO, group × time interaction, |t| > 2.1, cluster FWER corrected). **g)** fMRI connectivity between PFC and RS across time windows (↓Inhibition n=15; control n=17, group × time interaction, linear mixed-effects model). **h)** fMRI connectivity of the PFC in the window of maximal connectivity difference (15–35 minutes post CNO, group × time interaction, cluster FWER corrected, ↓Inhibition n=15, control n=17). BL: baseline; PFC, prefrontal cortex; RS, retrosplenial cortex; Th, thalamus; CPu, caudate–putamen. Normalized firing rate timeseries are shown as mean ± C.I. of loess fit. All other line plots show mean ± SEM; *p<0.05, ***p<0.001. Violin plots show median (thick line) and 75th and 25th percentiles (dashed line).

We first validated our chemogenetic manipulation by measuring electrophysiological activity in the PFC after administration of CNO in control and hM4Di-expressing mice (Figure 2). ↓Inhibition produced a sustained increase of neuronal firing in the PFC, consistent with heightened circuit-level excitability in the manipulated area (Figure 2b, p<0.001, Wilcoxon rank-sum, FDR corrected). This increase in MUA was accompanied by a prominent broadband increase in LFP power, spanning δ to high-γ frequencies (Figure 2c,d, cluster1=1.6-137 Hz, p<0.001, Student’s t-test, FWER-corrected). Thus, both ↑Excitation and ↓Inhibition increase local cortical excitability and firing rates, albeit with distinct electrophysiological oscillatory signatures. Importantly, as previously observed with ↑Excitation, ↓Inhibition also reduced mouse sociability as probed using a three-chamber sociability test (Figure S2c, p=0.034 Student’s t test). Together, these results document that ↓Inhibition is a physiologically effective and behaviorally relevant manipulation.

### Chemogenetic inhibition of PV+ interneurons results in fMRI hypoconnectivity

We next examined whether ↓Inhibition produces an fMRI signature comparable to that observed with ↑Excitation, despite the distinct LFP spectral profiles of the two manipulations. Consistent with our previous findings, **↓**Inhibition caused a significant BOLD signal increase in the PFC, partially extending into neighboring striatal regions (Figure 2e,f; p=0.023, group x time interaction, |t|>2.1, p<0.05 cluster-corrected). This increase in BOLD is consistent with the increase in high-frequency LFP power elicited by ↓Inhibition in the PFC (Figure 2b–d), and with the established coupling between high-frequency neuronal activity and local BOLD activity.

We next asked how ↓Inhibition affects long-range fMRI connectivity of the PFC. Seed-based mapping in DREADD-expressing mice revealed reduced fMRI connectivity between the PFC and some of its DMN targets, including the striatum, posterior cingulate, and retrosplenial cortex (Figure 2g,h, group x time interaction, |t|>2.1, p<0.05, FWER-corrected). Although weaker in magnitude and temporally shifted (peaking 15-35 min post-CNO; p=0.017, group x time interaction, Figure 2g, and S3b), the direction and anatomical distribution of the observed fMRI connectivity changes were broadly consistent with those elicited by ↑Excitation.

Notably, prior work suggests that DREADD-induced ↓Inhibition can lead to fMRI hypoconnectivity also in unimodal sensory cortices, such as the primary somatosensory cortex. Chemogenetic inhibition of PV+ interneurons in this region using a viral construct and sedation protocol analogous to those employed here was previously shown to reduce fMRI connectivity ^44^. To place these results within the same analytical framework of our PFC manipulations, we re-analyzed the corresponding fMRI data using our preprocessing steps and linear mixed-effects modelling. This analysis revealed reduced fMRI connectivity between the manipulated SSp and both contralateral and ipsilateral targets (Figure S6b, left, p = 0.012, group × time interaction; Figure S6b, right, |t|>2.1, p < 0.05, FWER-corrected). Together with our PFC results, these findings suggest that increased cortical excitability, whether driven by ↑Excitation or ↓Inhibition, is consistently associated with fMRI hypoconnectivity.

### Chemogenetic reduction of cortical excitability induces fMRI hyperconnectivity

To assess whether the inverse relationship between cortical excitability and fMRI connectivity is bidirectional, i.e., whether it also holds under conditions of reduced excitability, we re-analyzed a previously published dataset in which neuronal activity was chemogenetically suppressed in the mouse PFC ^16^. In that previous study, the inhibitory DREADD receptor hM4Di was expressed pan-neuronally in the PFC under the hSyn promoter, and CNO administration was used to suppress neuronal activity in the targeted region. Importantly, this manipulation robustly reduced spontaneous local firing activity ^16^, which we interpret here as reduced cortical excitability at the population level. We revisit here these prior data to relate their fMRI and electrophysiological responses to those produced by ↑Excitation and ↓Inhibition within a common analytical framework. For reference, we briefly recap these results in Figure 3 and denote this manipulation as “silencing”, in line with chemogenetic literature ^45^.

**Figure 3.**
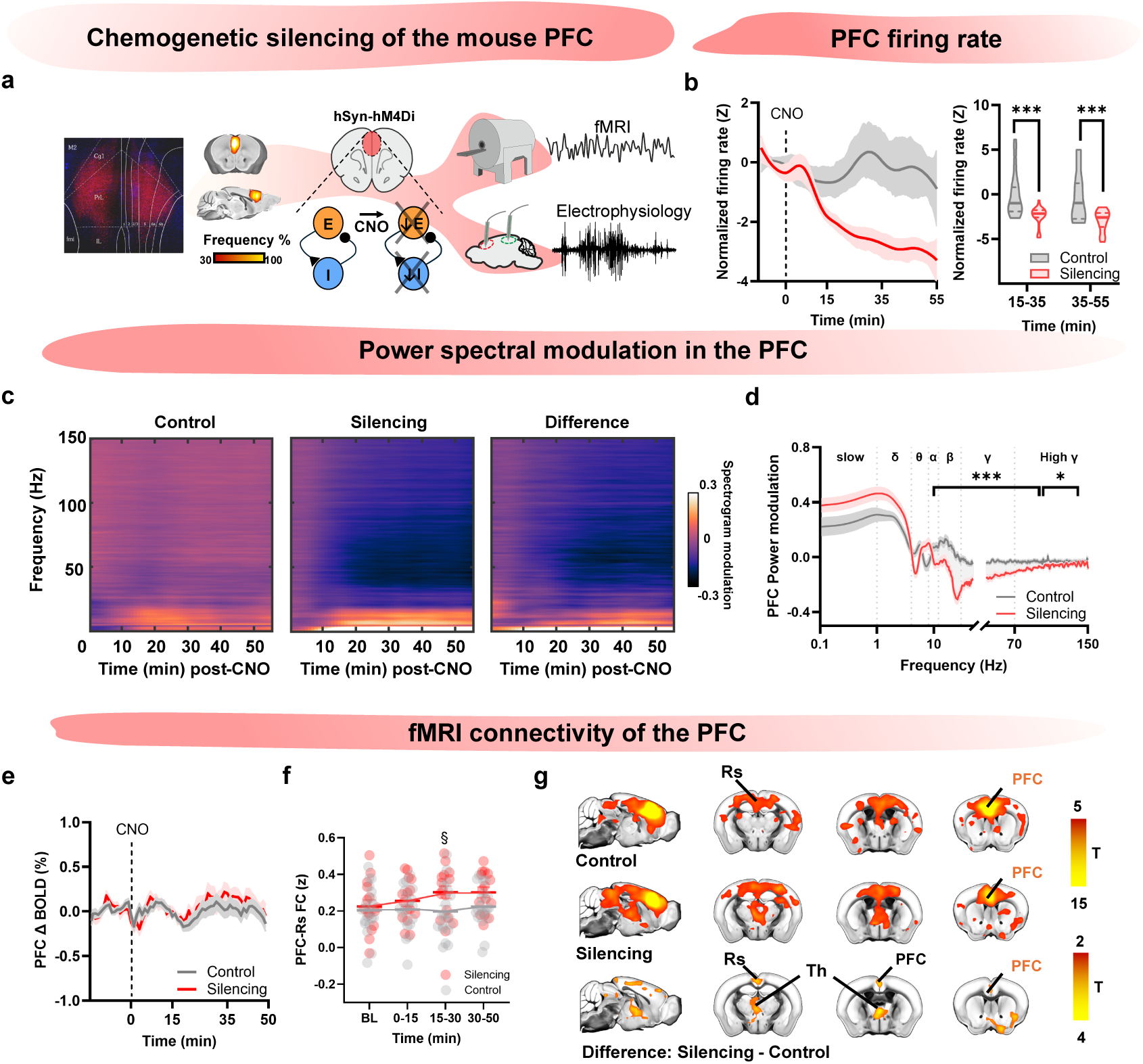
Chemogenetic silencing of the mouse PFC reduces local excitability and induces fMRI hyperconnectivity. **a)** Experimental design. hSyn-hM4Di was injected bilaterally in the medial PFC. Leftmost panels show representative viral expression and group-level viral coverage maps across animals. The effect of chemogenetic silencing was mapped using fMRI and multielectrode electrophysiology. **b)** Baseline-normalized firing rate in the PFC (right, summary over 15-35 and 35-55 min; Wilcoxon rank-sum test; n=100 time bins from n=5 control and n=5 Silencing mice). **c)** Group-average spectrogram modulation in PFC after CNO administration (control n=5; Silencing n=5), and difference (Silencing - control, right). **d)** Frequency resolved spectral modulation after chemogenetic silencing of the PFC (n=68 and n=60 bins from n=5 Silencing and n=4 control mice, respectively; two-sample t-tests, cluster FWER-corrected). **e)** Average fMRI BOLD signal change in PFC after CNO administration (Silencing n=15; control n=18). **f)** fMRI connectivity between PFC and RS across time windows (Silencing n=15; control n=18; two-sided Student’s t test). **g)** fMRI connectivity of the PFC 15–30 min post CNO and corresponding contrast map (Silencing > control; linear model followed by post hoc Tukey test; |t|>2.1, cluster FWER corrected). BL: baseline; PFC, prefrontal cortex; Rs, retrosplenial cortex; Th, thalamus; CPu, caudate– putamen. Normalized firing rate timeseries are shown as mean ± C.I. of loess fit. All other line plots show mean ± SEM. Timecourse plots show mean ± SEM; §p<0.05, uncorrected, *p<0.05, ***p<0.001. Violin plots show median (thick line) and 75th and 25th percentiles (dashed line).

Chemogenetic silencing of the PFC produced a marked reduction in firing rate in DREADD-expressing mice (Figure 3b). This effect was accompanied by a trend for increased slow (< 1Hz) and δ LFP power, together with a robust broadband decrease in α, β, γ and high-γ power in the manipulated region (Figure 3c,d; cluster1=9.8-100 Hz, p < 0.001; cluster2=101-129 Hz, p = 0.011; Student’s t-test). Quantification of BOLD fMRI signal in the manipulated region did not reveal significant BOLD signal changes (Figure 3e, p=0.395, group × time interaction). As reported in Rocchi et al., (2022), seed-based fMRI connectivity analysis revealed that chemogenetics silencing of the PFC increased fMRI connectivity between the manipulated area and multiple nodes of the mouse DMN, including thalamus, cingulate cortex, and retrosplenial cortices (Figure 3fg, general linear model, post-hoc Tukey test over 15-30 min post-CNO, |t|>2.1, p<0.05, cluster-corrected). Notably, in our prior study we also showed that this effect generalizes to different sedative mixtures (including the same sedation used for our other PFC studies), and to chronic virally-induced PFC silencing via potassium channel overexpression ^16^. Together with our ↑Excitation and ↓Inhibition results, these findings point to a bidirectional inverse relationship between cortical excitability and large-scale fMRI connectivity.

### Slow-band interareal coherence tracks fMRI connectivity across manipulations

Our results so far indicate that cortical manipulations that increase circuit-level excitability (↑Excitation and ↓Inhibition) lead to fMRI hypoconnectivity, whereas chemogenetic silencing of the PFC, a manipulation that reduces net circuit-level excitability, is associated with fMRI hyperconnectivity. How can local changes in cortical excitability bidirectionally shape large-scale fMRI connectivity? Because fMRI connectivity measures synchronous fMRI co-fluctuations (i.e., zero-lag correlation), we reasoned that the functional reconfiguration induced by our manipulations could be explained by changes in electrophysiological coupling across regions. To test this hypothesis, we simultaneously recorded LFP activity in the manipulated area and one of its main cortical anatomical projection targets (retrosplenial cortex, Rs) across all chemogenetic manipulations, and quantified magnitude-squared LFP coherence between electrodes in a post-injection time window identical to that used for fMRI mapping.

This analysis revealed different frequency-resolved interareal coherence profiles for each manipulation (Figure 4). ↑Excitation produced a robust reduction in PFC-Rs coherence that was most pronounced at low frequencies (slow i.e. <1 Hz, δ and θ-α), but also extended into the β range (Figure 4a, cluster1 = 0.1-0.9 Hz, p=0.044; cluster2 = 1.7-8.1 Hz, p < 0.001; cluster3 = 18.7-22 Hz, p = 0.011; Student’s t-test, FWER-corrected; Figure 4a, right, p < 0.001, Wilcoxon rank-sum test, FDR-corrected). ↓Inhibition yielded a composite coherence profile, characterized by decreased coupling at low frequencies (< 2 Hz), and increased coherence in faster rhythms (θ-β; Figure 4b, left; cluster1 = 0.1-1.7 Hz, p = 0.0116; cluster2 = 2.3-9.6 Hz, p < 0.001; cluster3 = 10-21 Hz, p < 0.001; cluster4 = 23-30 Hz, p < 0.001; Student’s t-test, FWER-corrected; Figure 4b, right, p < 0.001, Wilcoxon rank-sum test, FDR-corrected). By contrast, chemogenetic silencing increased coherence in slow (<0.1 Hz), δ and β bands. This effect was associated by a decrease in coherence in the α band (Figure 4c, cluster1=0.1-2.9 Hz, p < 0.001; cluster2=16-20.2 Hz, p=0.002; cluster3=32-34.3 Hz, p=0.029; Student’s t-test, FWER-corrected; Figure 4c, right, slow-δ p < 0.001, α p = 0.017, β p = 0.040, Wilcoxon rank-sum test, FDR-corrected). Together, these results show that changes in cortical excitability robustly alter interareal LFP coherence via dissociable, frequency-specific coupling profiles.

**Figure 4.**
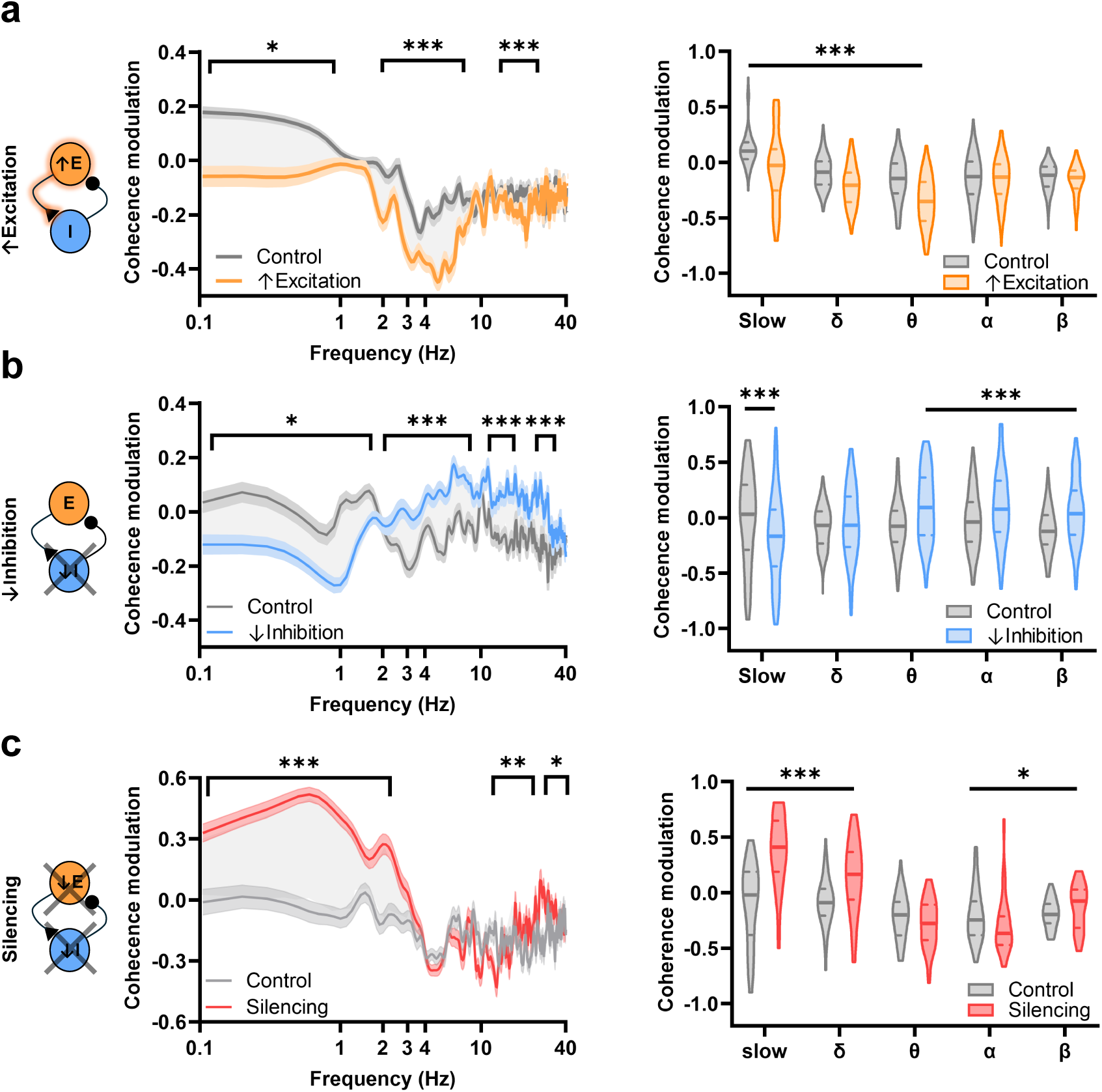
↑Excitation, ↓Inhibition and Silencing produce dissociable PFC-retrosplenial LFP coherence profiles. Frequency-resolved modulation of LFP coherence between PFC and retrosplenial cortex (Rs) (left) and corresponding band-limited summaries (right) for **(a)** ↑Excitation (↑Excitation, 120 bins, n=6 mice, control 160, n=7 mice) **(b)** ↓Inhibition (↓Inhibition 220 time bins, n=11 mice, control 200 bins, n=10 mice) and **(c)** Silencing (Silencing, 68 bins, n=5 mice, control 60 bins, n=4 mice). Frequency-resolved effects were assessed with two-sample t-tests and cluster-based FWER correction; band-limited effects were tested with two-sided Wilcoxon rank-sum tests with Benjamini–Hochberg FDR correction (q=0.05). Band-limited coherence modulation was computed for slow (0.1-1 Hz), δ (1-4 Hz), θ (4-8 Hz), α (8-12 Hz), and β (12-30 Hz) bands. Line plots show mean ± SEM; violin plots show median (thick line) and interquartile range (dashed lines). *p<0.05, ***p<0.001.

The divergence between these coherence profiles was particularly informative for ↑Excitation and ↓Inhibition, since both manipulations produced fMRI hypoconnectivity despite exerting dissociable effects on fast rhythms (i.e. θ-γ) coherence. Because spontaneous fMRI activity unfolds over very slow timescales, we hypothesized that fMRI connectivity would primarily reflect changes in low-frequency interareal coupling, irrespective of the different manipulation-specific coherency profiles at higher frequencies. This notion would be consistent with the observation that both ↑Excitation and ↓Inhibition reduce PFC-Rs coherence in low-frequency bands (Figure 4a,b), whereas chemogenetic silencing, which increases fMRI connectivity, enhances slow-band coherence (Figure 4c).

To test this hypothesis, we examined the covariance between fMRI connectivity and LFP coherence changes elicited by each chemogenetic manipulation across multiple frequency bands. We reasoned that electrophysiological coupling features relevant to fMRI connectivity should scale in magnitude with the corresponding fMRI connectivity changes across conditions. Consistent with this prediction, we found that among all tested frequency bands, low frequency coherence showed the strongest association with chemogenetically induced fMRI connectivity changes. Specifically, robust fMRI-LFP coherence correlations were observed within the low-frequency range (< 1 Hz) and across an extended 0.1-4 Hz range (Figure 5a and b, Spearman correlation, p=0.042 and p=0.006, respectively, FDR corrected). In contrast, coherence in higher-frequency bands showed weaker correlation, and non-significant associations with fMRI connectivity changes (Figure 5b). Because infraslow co-fluctuations of interareal γ-power (known as the γ-envelope) have also been proposed as a determinant of fMRI connectivity ^19^, we additionally tested whether coherence between the γ-power envelope timeseries in PFC and Rs correlated with the magnitude of our fMRI connectivity changes. This analysis did not reveal a significant association under our experimental conditions (Figure 5b, Spearman correlation ρ=0.50, p=0.216). Thus, among all electrophysiological measures tested, only slow (<4 Hz) interareal coherence systematically tracked fMRI connectivity across manipulations. This finding suggests that fMRI connectivity is primarily supported by, and preferentially sensitive to, low-frequency (<4 Hz) interareal electrophysiological coupling.

**Figure 5.**
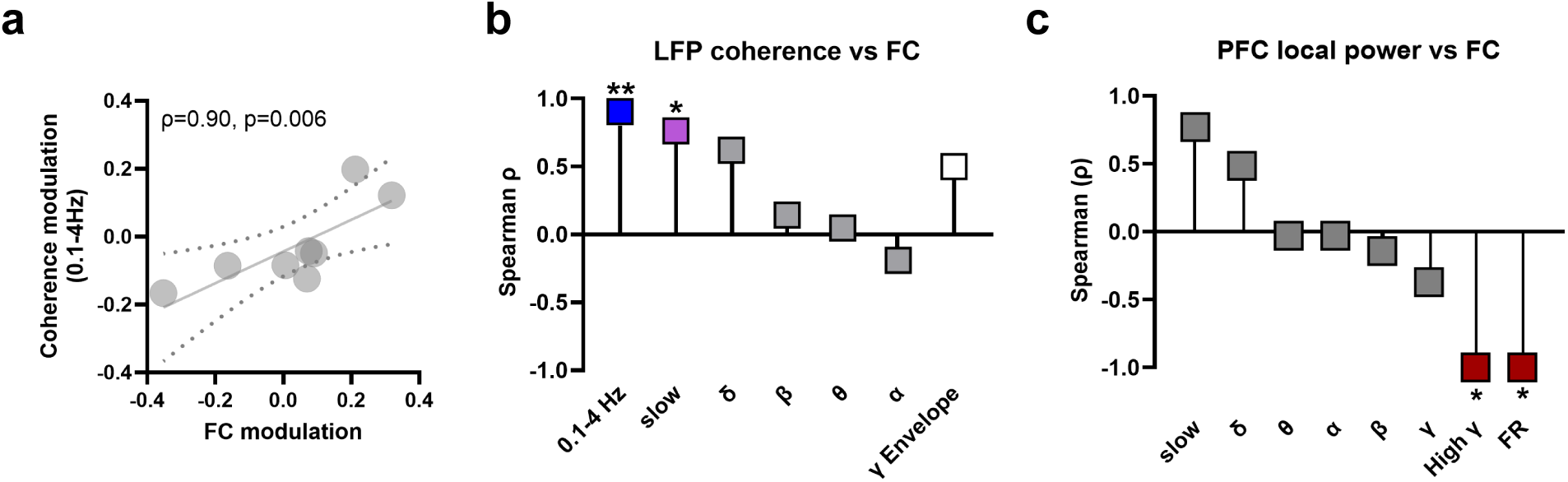
Cortical fMRI connectivity covaries with low-frequency LFP power coherence, and is inversely related to cortical excitability. **(a)** Correlation (Spearman) between fMRI connectivity modulation (FC) and LFP coherence modulation in the 0.1-4 Hz range. Dots indicate correlations between group-averaged FC and LFP coherence modulation across matched region-electrode pairs. **(b)** Band-resolved Spearman correlation between LFP coherence modulation and fMRI connectivity modulation (FC). (c) Band-resolved Spearman correlations between local PFC power modulation and fMRI connectivity modulation (FC; PFC-Rs and PFC-Th), including high-γ power as a proxy for local excitability. *p<0.05, **p<0.01, Benjamini-Hochberg corrected, q = 0.05. PFC, prefrontal cortex; Rs, retrosplenial cortex; Th, thalamus.

Cortical excitability is inversely related to fMRI connectivity

If low frequency interareal coherence provides a backbone for fMRI connectivity, how do local changes in circuit-level excitability influence this large-scale coupling? To address this question, we examined whether electrophysiological changes measured locally within the manipulated region alone (PFC) covaried with the corresponding differences in fMRI connectivity. We thus computed correlations between PFC-retrosplenial fMRI connectivity and average firing rate or band-limited LFP power modulation in the PFC across conditions. This analysis revealed a significant inverse correlation between fMRI connectivity and both average firing rate, and high-γ power-an established proxy of neuronal spiking activity ^46^ (Figure 5c, p=0.014, FDR corrected). This inverse relationship suggests that higher local excitability is associated with fMRI hypoconnectivity.

### A minimal biophysical model links cortical excitability, low-frequency coherence and large-scale coupling

Our data so far suggests that local cortical circuit-level excitability inversely modulates large-scale fMRI connectivity by altering low-frequency interareal neuronal coupling. We therefore considered a mechanistic account in which interareal coherence reflects at least two interacting components: a direct communication component, supported by high-frequency interareal electrophysiological coherence ^31, 47^, and a shared network-state component, driven by large-scale low-frequency covariations that synchronize distributed regions. Within this framework, fMRI connectivity primarily reflects the latter, low-frequency component.

To test this hypothesis, we built a minimal biophysical spiking network model in which a sending node (PFC; subjected to simulated chemogenetic manipulations) projects to a receiving node (RS) via direct excitatory connections (Fig. 6A). We also included a third node (a second sender region projecting to RS) that was not affected by the PFC manipulation, to model the fact that RS receives inputs from sources not influenced by the local PFC manipulation. Each node comprised recurrently interacting excitatory and inhibitory populations. In this model, direct communication between areas is implemented through firing-rate transmission, with oscillatory contributions to interareal coupling emerging only at higher frequencies (γ and high-γ coherence ^47^). However, all three areas also received a shared, slowly fluctuating input (with most power below 4 Hz) to capture large-scale state fluctuations. This configuration enabled us to examine how local E/I interactions, direct interareal connectivity, and global covariations jointly shape interareal coupling, and how these contributions are altered by chemogenetic perturbations.

**Figure 6.**
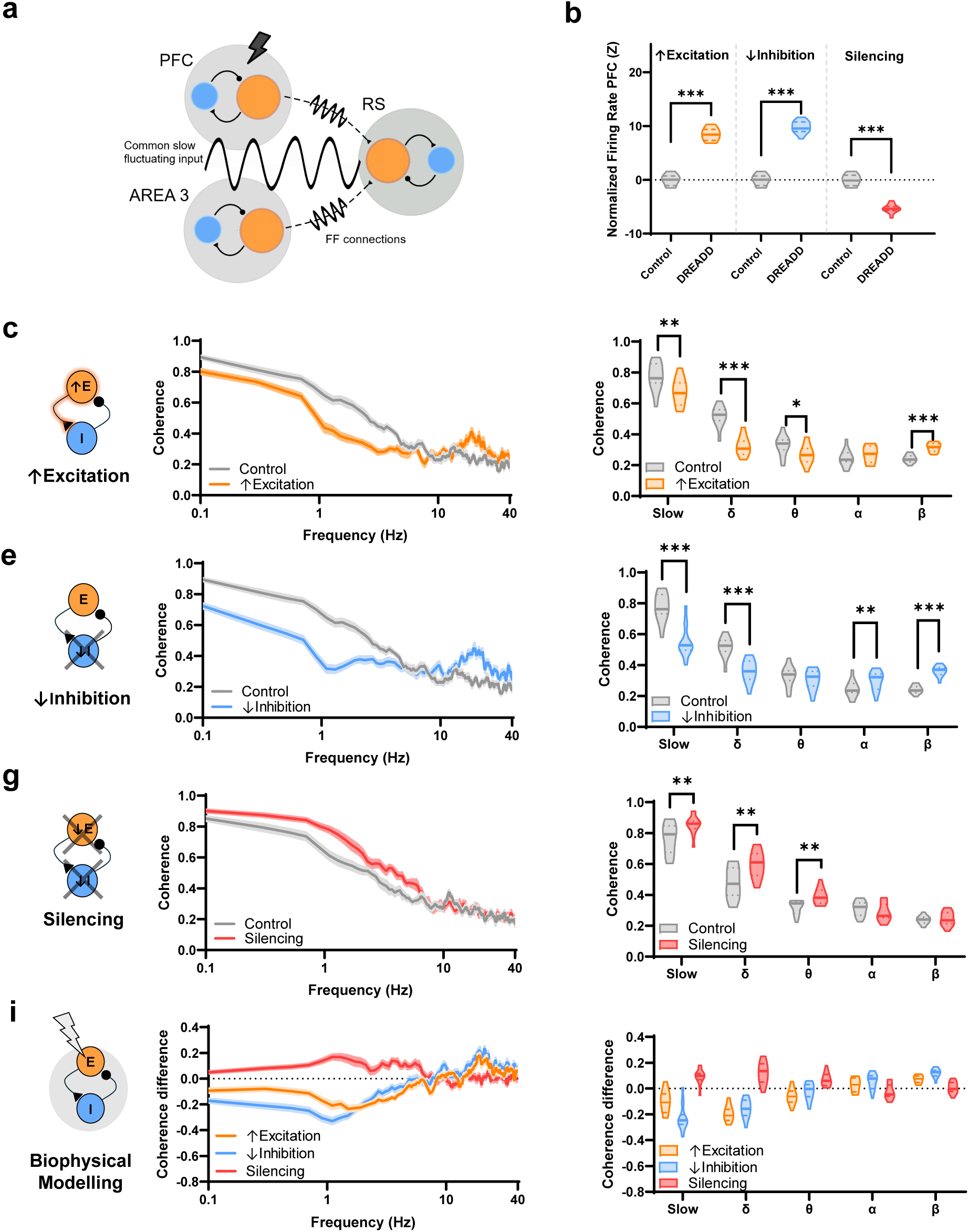
Simulated LFP coherence changes in a three-node network model reproduce empirical findings. **(a)** Model architecture: two recurrent excitatory-inhibitory (E-I) networks representing PFC and retrosplenial cortex (RS), were coupled via feedforward PFC→RS projections, and driven by a shared slowly fluctuating external input. A third node sending to RS models input to this region from remote regions not be affected by the PFC manipulation. **(b)** Simulated PFC firing-rate modulation under ↑Excitation, ↓Inhibition, and Silencing (normalized to control). **(c-h)** PFC-RS magnitude-squared (M-S) coherence in control and manipulation simulations for ↑Excitation **c-d)**, ↓Inhibition **(e-f)**, and Silencing **(g-h)**, shown as frequency-resolved spectra (left) and band-averaged coherence (right; slow 0.1-1 Hz, δ 1-4 Hz, θ 4-8 Hz, α 8-12 Hz, β 12-30 Hz). **(i-j)** Coherence difference, computed as the coherence in each experimental trial minus the average coherence across control trials. Line plots show mean ± SEM; violin plots show median and interquartile range.

Using this architecture, we mimicked the three chemogenetic manipulations in the sending area by systematically adjusting intrinsic single-neuron excitability and recurrent synaptic coupling in the “PFC” node (i.e., the manipulated site) through changes in resting membrane potential and synaptic conductance that mimic the reported effects of the different types of DREADD ^25^. Model-derived LFP signals were computed from the population synaptic activity in each area ^48^, and interareal coupling was quantified analogously to the empirical data as the magnitude-squared coherence between simulated PFC and RS LFPs. This minimal parameterization allowed us to ask whether changes in local circuit excitability alone, on top of the fixed anatomical coupling between areas, are sufficient to reproduce the frequency-specific reconfiguration of interareal coupling observed experimentally.

We first asked whether the parameters adjustments selected to mimic our chemogenetic manipulations (see Methods and Bertelsen, Mancini ^30^) reproduced the firing rate changes observed *in-vivo.* In line with our experimental recordings, simulations with parameters used to mimic ↑Excitation or ↓Inhibition produced a significant increase in PFC firing rate, whereas simulations with parameters chosen to mimic Silencing robustly suppressed neuronal firing (Figure 6b, p<0.001, Wilcoxon rank-sum test, FDR corrected). Thus, parameter settings chosen to reflect the known effects of each chemogenetic manipulation recapitulated the direction of firing-rate changes observed *in vivo*, thereby supporting the physiological plausibility of the model.

We next asked whether the model could reproduce the changes in interareal LFP coherence profiles elicited by each manipulation. Simulation of ↑Excitation reduced PFC-RS coherence in the slow-θ bands (Figure 6c,d, 0.1-1 Hz, p=0.007, δ, p<0.001, θ, p=0.029, Wilcoxon rank-sum test, FDR corrected), while increasing coherence in the β band (Figure 6c,d, p<0.001, Wilcoxon rank-sum test, FDR corrected). Simulation of ↓Inhibition produced a related pattern, with reduced low-frequency band coherence (0.01-1 Hz, p<0.001, δ, p<0.001, Wilcoxon rank-sum test, FDR corrected) and robustly increased coherence in α and β bands (α, p=0.008, β p<0.001, Wilcoxon rank-sum test, FDR corrected, Figure 6e,f). Thus, in our *in-silico m*odel, manipulations that enhance local excitability consistently weakened low-frequency interareal coherence. In contrast, simulated PFC silencing produced the opposite effect, increasing coherence in the slow (0.1-1 Hz) and δ bands (p=0.006 and p=0.003 respectively), as well as in the θ band (p=0.002, Wilcoxon rank-sum test, FDR corrected, Figure 6g,h).

Although our model did not capture the entire range of all the spectral features we measured experimentally, it crucially reproduced the direction of low-frequency coherence changes, i.e., the electrophysiological measure that best tracked changes in fMRI connectivity produced by all three manipulations. Specifically, ↑Excitation and ↓Inhibition both reduced low-frequency coherence in the slow and δ bands, whereas Silencing increased it (Figure 6i,j). Together, these results provide a mechanistic account of how cortical excitability can shape large-scale fMRI network synchronization: recapitulating our empirical data, they indicate that local changes in cortical excitability are sufficient to modulate slow-band interareal coupling, and that bidirectional changes in excitability shape fMRI connectivity primarily by locally interfering with slow-band electrophysiological coherence.

Collectively, the findings we describe in the present work motivate a simple two-pillar conceptual framework for understanding the neuronal rhythms underlying large-scale fMRI connectivity and their modulation by intrinsic cortical excitability (Figure 7). The first pillar posits that low-frequency (slow and δ bands) interareal coherence provides the electrophysiological scaffold for large-scale fMRI connectivity. The second pillar asserts that local cortical excitability is inversely related to fMRI connectivity. Within this framework, regions within a functional network are synchronized primarily via spatially distributed, coherent low-frequency rhythms. Elevating local circuit-level excitability increases high-frequency neuronal activity and biases interareal communication towards direct area-to-area coupling, thereby reducing the contribution of coherent low-frequency interareal fluctuations and weakening large-scale fMRI coupling of the affected region. Conversely, reducing excitability facilitates the synchronization of low-frequency rhythms, thereby enhancing network integration and fMRI connectivity. This framework parsimoniously explains the seemingly counterintuitive effects of excitability changes reported here and elsewhere, and suggests that higher-frequency rhythms influence fMRI connectivity mainly through their interaction with slow oscillatory activity.

**Figure 7.**
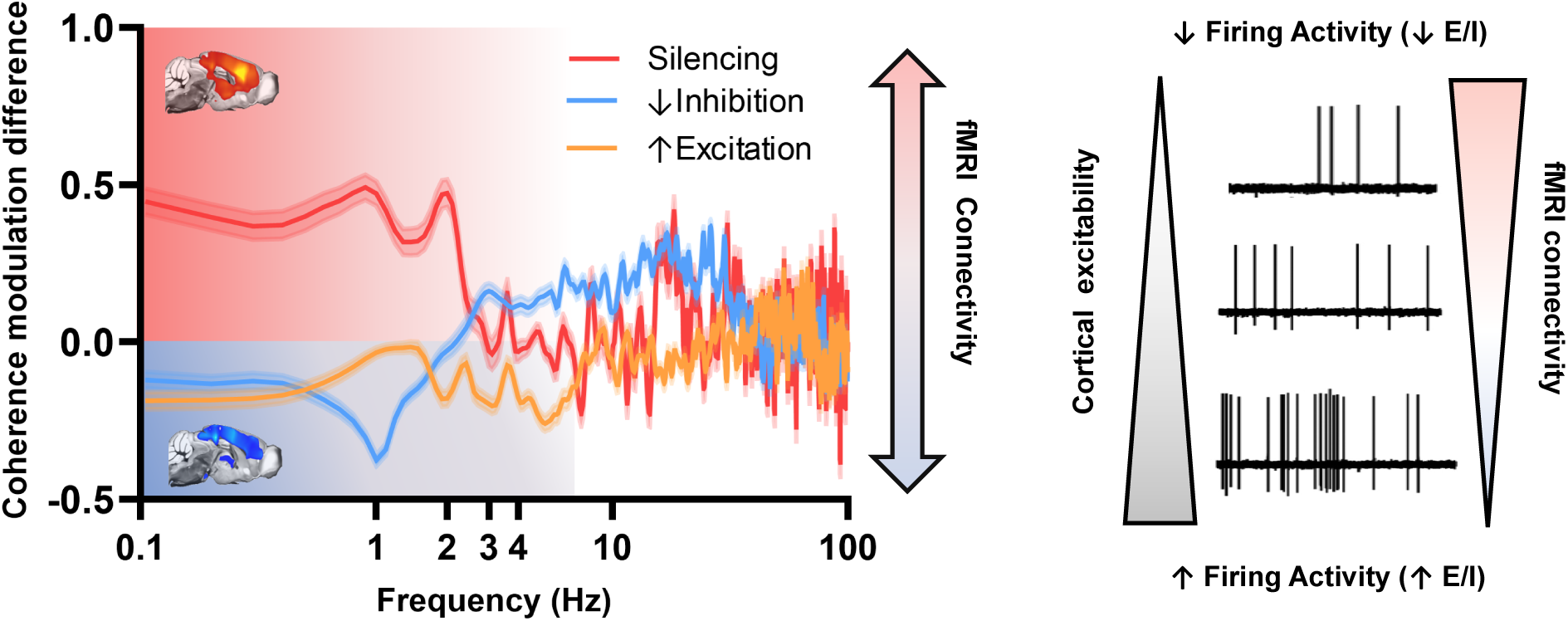
A mechanistic framework linking cortical excitability, low-frequency interareal coherence, and fMRI connectivity. Low-frequency interareal coherence provides the electrophysiological scaffold for large-scale fMRI connectivity. Changes in local cortical excitability (indexed by firing activity) modulate network coupling by weakening or facilitating this low-frequency synchrony: increased excitability weakens slow coherent fluctuations (here illustrated as coherency modulation difference) and reduces fMRI connectivity, whereas reduced excitability promotes slow synchrony and enhances fMRI connectivity.

## Discussion

In the present study, we aggregated cell type-specific chemogenetic perturbations that either increase or reduce cortical excitability and examined their impact on large-scale fMRI connectivity and electrophysiological coupling. Using this approach, we found that cortical excitability, operationalized as regional firing activity, is inversely related to large-scale fMRI connectivity. Importantly, by comparing these perturbations within a unified analytical framework, we also found that only low-frequency (<4 Hz) interareal coherence systematically tracks the direction and magnitude of the ensuing fMRI connectivity changes. Together, our results identify cortical excitability as a latent systems-level physiological variable that robustly shapes macroscale fMRI connectivity, and highlight low-frequency interareal coherence as a key physiological correlate of this coupling.

The results of the present study converge onto a two-pillar framework that has potential far-reaching implications for the interpretation of fMRI connectivity as a readout of interareal synchronization and regional cross-talk. The first tenet of this framework is that regional cortical excitability is inversely related to large-scale fMRI connectivity, i.e., regions that are in a hyperexcitable state show reduced fMRI connectivity, whereas regions that are in a less excitable, electrophysiologically quieter state, show increased fMRI connectivity. These observations challenge the often-implicit notion of a monotonic relationship between a region’s metabolic or neuronal activity, and its functional connectivity^49^, and instead support a view in which node-specific shifts in cortical excitability are sufficient to robustly shape and reconfigure macroscale network coupling.

Multiple theoretical and experimental studies are consistent with this framework. For example, in good agreement with our empirical results, recent whole-brain modelling in the mouse showed that the effect of regional brain perturbations on fMRI connectivity is inversely related to the estimated firing-rate changes produced by the employed manipulations^27^. Cortical desynchronization consistent with the fMRI hypoconnectivity we describe here has also been recently described in ↓Inhibition chemogenetic experiments conducted in awake mice using widefield calcium imaging^50^. Human studies also support our framework. For example, EEG and fMRI recordings in hemispherectomy or brain-injured patients shows that fMRI hyperconnectivity can reflect sleep-like, diffuse low-frequency synchronization, and that preserved network patterns can coexist with electrophysiological signatures of cortical silencing^51, 52^. Similarly, bidirectional changes in functional connectivity mirroring the effects described here have been reported in neurostimulation studies in which motor or prefrontal cortical regions are inhibited or excited using TMS stimulations at different frequencies^53, 54, 55^.

The presence of an inverse relationship between excitability and functional connectivity may be of particular relevance for brain disorders linked to alterations in cortical excitability such as autism^35^, or schizophrenia^56, 57^. Human studies have revealed a heterogeneous landscape of fMRI connectivity alterations (dysconnectivity) in these disorders, spanning both hyper- and hypoconnectivity even within seemingly homogeneous populations^58, 59, 60, 61^. This variability is mirrored by preclinical models^62, 63^. Our data suggest that divergent patterns of fMRI dysconnectivity may partly reflect bidirectional node-specific in cortical excitability. In keeping with this, recent EEG studies using the Hurst exponent (a metric sensitive to E/I balance^30, 36^ show that autistic children can be stratified into dominant subgroups with higher or lower cortical excitability, linked to distinct behavioral outcomes^30^. It is thus possible that these two diverging EEG subtypes could be associated with hypo- and hyperconnectivity, respectively, thus revealing a plausible electrophysiological substrate for dominant fMRI neurosubtypes identified in autism^63^. Further corroborating this framework, fMRI connectivity in autism has been shown to inversely track with EEG-based measures of cortical excitability^36^, and investigations in autistic individuals have linked decreased cortical inhibition with increased functional connectivity^64^. More broadly, our findings support the notion that atypical fMRI connectivity and E/I imbalance are causally interrelated, explaining why disorders characterized by alterations in cortical excitability very often show also abnormal fMRI connectivity^60, 65, 66^.

Importantly, an inverse relationship between excitability and connectivity also offers a putative parsimonious explanation for seemingly counterintuitive reports of fMRI hyperconnectivity in neurodegenerative or lesional conditions. Within our framework, such findings can be interpreted as a direct consequence of reduced spontaneous firing in partially deafferented or cell-loss-affected regions, which might favor slow, hypersynchronous network dynamics^52, 67^, rather than (or in addition to) compensatory network reorganization^68^. We finally note that this relationship may also partly explain developmental changes in fMRI connectivity related to the maturation of inhibitory systems^69^. In keeping with this, it has been recently shown that a transient increase in cortical excitability during developmental can permanently increase cortical excitability, resulting in autism-relevant hypoconnectivity ^70^. These findings are consistent with cortical excitability acting as a systems-level physiological axis that bidirectionally controls fMRI connectivity in disease states, as well as during healthy developmental and aging.

Our experimental results, corroborated by our minimal biophysical model, also point to a key contribution of low-frequency (< 4Hz) fluctuations as a neuronal backbone for interareal fMRI connectivity. This second conceptual pillar substantiates and extends previous investigations of the neurophysiological basis of fMRI connectivity, corroborating the dominance of slow/infraslow rhythms in sustaining spontaneous fMRI activity^12, 13, 15, 20, 71, 72^, and suggesting that electrophysiological coherence in low-frequency, but not faster rhythms, primarily supports fMRI connectivity. An intriguing postulate of our findings is that increased coherence in higher frequency rhythms (as observed under ↓Inhibition), does not necessarily result in fMRI hyperconnectivity, no matter how big the corresponding coherence changes are. This observation marks a substantial difference between hemodynamic (fMRI-like) and electrophysiological measures of functional connectivity, suggesting that, despite its richer spatial resolution, fMRI connectivity is disproportionately sensitive to a limited portion of the full electrophysiological spectrum. This distinction should thus be accounted for when using fMRI connectivity to investigate the online effect of high-frequency neurostimulation or fast-oscillation entrainment^53, 73, 74, 75^.

We finally note three considerations that delimit the scope of the present framework. First, while our aggregate analyses support an inverse relationship between local excitability and fMRI connectivity, not all chemo-fMRI studies map directly onto our experimental manipulations ^27, 76^. Because ostensibly similar chemogenetic manipulations can place neocortical circuits in different effective physiological regimes^26, 28^, and given that the population-level effects of chemogenetic manipulations are rarely quantified or validated *in vivo*, the physiological basis of these discrepancies remains to be determined. Pairing fMRI with direct measurements of population activity and interareal coupling will thus be essential to determine whether apparent deviations from our model reflect genuine mechanistic divergence, or instead arise from different effective cortical regimes induced by specific perturbations. Second, our experiments and modelling focus mostly on corticocortical interactions. It thus remains an open question whether (and to what extent) the same principles apply to subcortical structures, where local circuit motifs, long-range drive, and dominant coupling frequencies can differ markedly. Third, while cortical excitability provides a compact explanatory axis for the direction of the connectivity changes, our work does not imply that this is the only neurophysiological mechanism affecting fMRI connectivity and its alterations. Neuromodulatory tone^77, 78, 79^ and arousal state^79, 80^, for example, are known to shape local gain and cortical activity, thus influencing the direction of fMRI connectivity both directly and potentially by altering cortical E/I balance^81^. Our findings suggest that regional alteration in cortical excitability, irrespective of their primary physiological origin, can robustly influence large-scale fMRI connectivity on top of many other, potentially more nuanced, mechanisms, including purely neurovascular contributions^82^.

In summary, by integrating cell type-specific chemogenetic perturbations with multielectrode physiology in a unified analytical framework, we identify an inverse relationship between regional cortical excitability and large-scale fMRI connectivity, and show that the direction and magnitude of these connectivity changes are tracked by slow-band (<4 Hz) interareal coherence. These results shed light on the neural correlates of large-scale fMRI connectivity and provide a parsimonious mechanistic framework for interpreting dysconnectivity patterns linked to altered cortical excitability.

## Methods

### Ethical statement

All *in in vivo* experiments were conducted in accordance with the Italian law (DL 26/2014, implementing EU directive 2010/63), and were approved by the Animal Care Committees of the University of Trento, the Istituto Italiano di Tecnologia, and the Italian Ministry of Health. All surgical procedures were performed under deep anesthesia.

### Animals

CamkII-hM3Dq-mediated excitation (↑Excitation) and hSyn-hM4Di pan-neuronal inhibition (Silencing) experiments were carried out in adult C57BL/6J mice (Jackson Laboratory, stock no. 000664). hM4Di-mediated inhibition of parvalbumin-positive interneurons (↓Inhibition) was carried out in a transgenic PV-Cre line, (B6;129P2-Pvalb^tm1(cre)Arbr^/J; Jackson Laboratory, stock no. 017320), in which Cre recombinase is expressed under the parvalbumin promoter. Mice were housed in groups under a 12:12 h light–dark cycle with food and water available ad libitum. Environmental conditions were maintained at 21 ± 1 °C and 60 ± 10% relative humidity.

### Experimental cohorts

This study aggregates both new experimental studies and previously published datasets. All datasets were acquired by the Gozzi lab at the IIT unless otherwise stated. Experimental cohorts for newly generated datasets were composed as follows.

#### Behavioral testing

A social interaction test was performed in four separate cohorts of adult male mice (9.7 ± 4.8 weeks at the age of surgery): ↑Excitation control (n=31); ↑Excitation (n=31); ↓Inhibition control (n=14); ↓Inhibition (n=18).

#### Chemo-fMRI

fMRI acquisitions in sedated animals were performed in four separate cohorts of adult male mice (11.3 ± 7.8 weeks at the age of surgery): ↑Excitation control (n = 17), ↑Excitation (n = 17), ↓Inhibition control (n = 17), and ↓Inhibition (n = 15). Awake fMRI recordings were performed in a separate cohort of adult male mice (35± 4.8 weeks at the age of surgery): ↑Excitation (n = 10) and control (n = 14).

#### Electrophysiology

Multi-electrode recordings in the ↑Excitation and ↓Inhibition studies were conducted in adult male mice (11 ± 4.80 weeks at the age of surgery) using the following cohorts: ↑Excitation control (n = 7), ↑Excitation (n = 6), ↓Inhibition control (n = 10), and ↓Inhibition (n = 11). One ↑Excitation mouse was excluded from the multi-unit activity (MUA) analysis due to unreliable spike detection throughout the recording but was retained for LFP analyses. Chemogenetic activation in the ↑Excitation and ↓Inhibition experiments employed different CNO doses (↑Excitation: 0.5 mg/kg, s.c.; ↓Inhibition: 2 mg/kg, i.v.), reflecting the different affinities of hM3Dq and hM4Di receptors. To control for potential off-target effects of CNO, the drug was administered to both control and experimental groups so that any inter-group difference would primarily reflect DREADD receptor activation.

The following previously published datasets were also used in this study.

#### Chemo-fMRI

Chemogenetic silencing of the PFC was described in Rocchi, Canella ^16^ and was performed in adult male mice (10.7 ± 2.3 weeks at the age of surgery): control (n = 19) and Silencing (n = 15).

fMRI acquisitions in mice undergoing chemogenetic modulation of excitability in somatosensory cortex were previously described in Markicevic et al. ^44^ and were acquired at ETH (Zurich). Experimental cohorts consisted of adult male C57BL/6J mice (9.9 ± 1.6 weeks at the age of surgery) and included a single control group (n = 13), which served as the reference for both the ↑Excitation cohort (n = 13) and the ↓Inhibition cohort (n = 19; Pvalb^tm1(cre)^Arbr, PV-Cre mice). DREADD receptor activation in these studies was induced by intravenous clozapine administration (30 μg/kg) to both experimental and control cohorts 15 minutes after the beginning of the scan^78^.

#### PCP modulation of fMRI connectivity

fMRI recordings examining PCP-induced modulation of connectivity were previously described in Montani et al., ^41^. Recordings were performed in adult male C57BL/6J mice (12 weeks old) receiving either PCP (n = 10) or vehicle (saline; n = 10). PCP was administered intravenously at 0.5 mg/kg.

#### Electrophysiology

In this study we reanalyzed both single-electrode and multielectrode recordings from Rocchi et al., (2022). Chemogenetic silencing of the prefrontal cortex (PFC) was performed in adult male mice (18-30 weeks old at the time of surgery). Single-electrode recordings were used to quantify PFC firing rates. In this study experimental cohorts consisted of a silencing group (n = 5) and a control group (n = 5), with recordings lasting 55 minutes following CNO administration. For multielectrode local field potential (LFP) recordings, cohorts consisted of a silencing group (n = 5) and a control group (n = 4), with recordings lasting 30 minutes following CNO administration. CNO was administered intravenously to both control and silencing cohorts at a dose of 2 mg/kg.

### Viral injections

Before surgery, mice were anesthetized with isoflurane (4% induction, 2% maintenance) and placed in a stereotaxic apparatus (Kopf). Injections were performed with a Hamilton syringe mounted on Nanoliter Syringe Pump with controller (KD Scientific), at a rate of 0.08 nl/s, followed by a 15 min waiting period to minimize backflow. Viral injections targeted the medial prefrontal cortex (PFC), here defined as a meta-region including the anterior cingulate, prelimbic and infralimbic cortices^83^, corresponding to a key hub of the mouse default mode network^84^. All viral transductions were carried out using high-titer viral suspensions (>10^13^ vg/mL).

To chemogenetically stimulate principal neurons in the PFC (↑Excitation), 250 nl of AAV9-CaMKIIa-hM3D(Gq)-mCherry (Addgene, #50476-AAV9, titer≥1×10¹³ vg/mL) were injected bilaterally into the PFC (total volume of 500 nL; coordinates from Bregma: AP +1.7mm, ML +/- 0.3 mm, DV 1.7 mm) as previously described ^16^. To inhibit PV^+^ interneurons (↓Inhibition), 400 nL of AAV8-hSyn-DIO-hM4D(Gi)-mCherry (Addgene, #44362-AAV8 titer ≥ 1×10¹³ vg/mL) were injected bilaterally into the PFC of PV-Cre transgenic mice (total volume, 800 nL). Sham surgeries were performed in control mice.

Injection of the hM4Di DREADD receptor used for pan-neuronal silencing of the PFC has been originally described in Rocchi, Canella ^16^. Briefly, AAV8-hSyn-hM4D(Gi)-mCherry was injected bilaterally (1 µL per hemisphere; total volume 2 µL) at the same coordinates used for the ↑Excitation and ↓Inhibition experiments. Viral transductions were carried out using high-titer viral suspensions (>10^13^ vg/mL).

Chemogenetic manipulation of cortical excitability in the right primary somatosensory cortex (SSp) was carried out at ETH Zurich as previously described^44^. Briefly, to chemogenetically activate principal neurons in the right SSp (↑Excitation), 950 nL of ssAAV8-mCAMKIIα-hM3D-mCherry (titer≥1.8 × 10^12^ vg/mL) were injected unilaterally at the following stereotaxic coordinates (Coordinates from Bregma: AP +0.5 mm, ML-3.0 mm, DV-2.25 mm). To inhibit PV^+^ activity in the right SSp, 950 nL of ssAAV8-hSyn1-dlox-hM4Di-mCherry(rev)-dlox-WPRE-hGHp(A) (titer≥5.4 × 10^12^ vg/mL) were injected at the same stereotaxic coordinates.

### Behavioral tests

Sociability was probed using the three-chamber test as previously described ^70^. Stimulus mice were experimentally naïve and matched to experimental mice in sex, genetic background, age, and weight. Four days prior to testing, stimulus mice were habituated to the wire cups. The test began with a 10-min habituation to the central compartment, followed by a 10-min habituation to all three chambers. Immediately thereafter, the 10-min sociability phase commenced, during which the experimental mouse was placed in the central compartment and allowed to explore the lateral chambers containing either a stimulus mouse or an empty cup. The positions of the stimulus and empty cups were randomized across trials. Clozapine-N-oxide (CNO, Sigma Aldrich) was administered via intraperitoneal injection 30 min before the first habituation phase in all experimental and control subjects. No overt behavioral signs of epileptic seizures were observed in any animal during testing. To quantify the effect of ↑Excitation and ↓Inhibition on social behavior, we calculated a sociability index (SI) as previously described^85^. The SI accounts for the time spent investigating the social stimulus (T_social_) versus the empty cup (empty cup, T_control_), and was computed as follows:

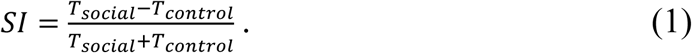

### fMRI acquisitions

↑Excitation and ↓Inhibition fMRI scans were acquired as previously described ^16^. Mice were intubated and tail-vein cannulated under isoflurane anesthesia (2%) prior to scanning. Following positioning in the scanner, sedation was induced with a low-dose combination of medetomidine and isoflurane (med-iso). An initial intravenous bolus of medetomidine (0.05 mg/kg) was administered, followed 5 min later by a continuous infusion (0.1 mg/kg/h) with isoflurane maintained at ∼0.5%. fMRI acquisition began 30 min after med-iso induction.

Functional images were acquired with a 7 T MRI scanner (Bruker, Ettlingen) equipped with a BGA-9 gradient set (380 mT/m, max. linear slew rate 3,420 T/m/s), using a 72 mm birdcage transmit coil and a 4-channel solenoid coil for signal reception. The scanner was operated using Paravision 6.01 software (Bruker, Ettlingen). fMRI time series were acquired using single-shot echo-planar imaging with the following parameters: TR/TE, 1000/15 ms; flip angle, 60°; matrix, 98 × 98; field of view, 2.3 × 2.3 cm; 18 coronal slices; slice thickness, 550 µm; bandwidth, 250 kHz.

For all experiments, two consecutive 37-min acquisitions were carried out. The first 2 min (120 volumes) of each scan were excluded from analysis to allow for gradient equilibration, yielding two 35-min time series (2,100 volumes each) per subject. CNO doses were selected based on pilot experiments and previous reports^16, 24, 86^. CNO was administered intraperitoneally in both experimental and control animals (↑Excitation experiment=0.5 mg/kg, ↓Inhibition experiment=2 mg/kg) 17 min after the start of the first acquisition, yielding a 15-minute (900 volumes) pre-injection window that we used as baseline reference.

Awake fMRI acquisitions were performed following the surgical and habituation procedures previously described in ^42^. Briefly, mice were implanted with a custom-made headpost cemented to the skull. After a recovery period, mice underwent a two-steps habituation protocol spanning 26 days. During the first step (days 1-8), mice were acclimated to handling for 15 minutes per day over 3 consecutive days, followed by two days (days 4-5) of 10-minute cradle exploration sessions during which brief manual head fixation was introduced by holding the headpost for a few seconds. The second step (days 8-26) consisted of habituation to a mock scanner environment. During these sessions, the headpost was secured to the cradle; mice were initially left unrestrained (days 8-19) and were progressively introduced to gentle body restraint by taping them to the cradle toward the end of the protocol (days 22-26), allowing normal breathing and minimal movement. Throughout mock scanning, audio recordings of EPI pulse sequences were played at sound noise levels matching those of the actual MRI acquisition.

To minimize stress associated with prolonged awake restraint, and to match the experimental timeline we previously used to reliably map fMRI networks in awake mice ^42^, we employed a shorter acquisition session than in sedation studies. Specifically, each session consisted of a single 40-min acquisition comprising a 10-min pre-CNO baseline scan (600 volumes) immediately followed by a 30-min post-CNO scan (1,800 volumes), acquired using the same parameters as the med-iso dataset. Clozapine-N-oxide (CNO) was administered intraperitoneally (0.5 mg/kg).

Below, we also provide a brief summary of the experimental conditions for the previously published datasets used in this study. Chemo-fMRI acquisitions of PFC silencing were carried out at IIT and originally described in Rocchi et al.,^16^. Two consecutive fMRI scans (35 min, 2100 volumes; followed by 30 min, 1800 volumes) were acquired under halothane sedation. Animals were initially anesthetized with isoflurane (4% induction, 2% maintenance), intubated, and intravenously-cannulated for subsequent CNO administration. After positioning the mouse in the scanner, isoflurane was discontinued and replaced with halothane sedation (0.8%). fMRI acquisition began ≥30 min after isoflurane cessation to allow anesthetic washout. CNO (2 mg/kg) was administered intravenously 15 min after the start of the first acquisition. In the same study we showed that similar fMRI effects are obtained using med-iso sedation.

Pharmacological fMRI acquisitions with phencyclidine (PCP) were originally described in Montani et al.,^41^. In this study, mice were anesthetized with isoflurane (4%), intubated, and artificially ventilated. Forty-five minutes before acquisition, isoflurane was replaced with halothane (0.8%), and scans were performed under halothane sedation (0.8%). A 30-min fMRI scan was acquired, and 10 min after scan onset, either vehicle (saline, n = 10) or PCP (1 mg/kg, n = 10) was administered intravenously. BOLD acquisitions were performed with single-shot echo-planar imaging using parameters different from the previous datasets: TR/TE, 1,200/15 ms; flip angle, 30°; matrix, 100 × 100; field of view, 2 × 2 cm; 18 coronal slices; slice thickness, 600 µm.

fMRI acquisitions during chemogenetic manipulations of the mouse SSp were originally described in Markicevic at al., ^44^. Experiments were carried out at ETH (Zurich) on a 7-T Bruker BioSpec using receive-only cryogenic coil (Bruker BioSpin AG, Fällanden, Switzerland), combined with a linearly polarized room temperature resonator for RF transmission. Imaging parameters were: TR/TE, 1000/ 15 ms; flip angle, 60°; matrix size, 90 × 50; in-plane resolution, 0.2 × 0.2 mm^2^; 20 coronal slices; slice thickness, 400 µm. Mice were anesthetized with isoflurane (4%), intubated, vein-cannulated and mechanically ventilated. After positioning, a bolus of medetomidine (0.05 mg/kg) and pancuronium (0.25 mg/ kg) was administered intravenously and isoflurane reduced to 1%. Five minutes later, continuous infusion of medetomidine (0.1 mg/kg/h) and pancuronium (0.25 mg/kg/h) was initiated and isoflurane further reduced to 0.5%. ↑Excitation mice were scanned for 60 min (3600 volumes) and ↓Inhibition mice for 45 min (2700 volumes). Clozapine (30 µg/kg) was intravenously injected 15 min after acquisition onset in all groups.

### Image preprocessing and analysis

#### BOLD response

To map BOLD signal intensity changes in the PFC due to our manipulations, fMRI time series were pre-processed as follows. The first 120 volumes of each timeseries were discarded removed to account for gradient thermal equilibration. Data were then despiked, motion corrected, and co-registered to an in-house functional reference template. To preserve slowly evolving fMRI signal changes produced by CNO, no nuisance regression or band-pass filtering was applied. After spatial normalization, the two EPI timeseries of each animal were concatenated. To correct for intensity differences due to RF auto-equilibration between acquisitions, the difference between the mean signal intensity of the last 30 volumes of the first scan and the first 30 volumes of the second scan was subtracted from the second time series. Fractional BOLD signal intensity (%BOLD) was then computed voxel-wise from the concatenated time series relative to the pre-CNO baseline (ΔBOLD). To correct for nonlinear baseline trends present in our recorded BOLD signal and scanner drift, we first computed a voxel-wise mean time course across control animals and fit each voxel’s mean signal with a third-order polynomial (MATLAB polyfit), after normalizing the time axis to −1 to 1 to improve numerical stability. These voxel-wise polynomial fits were interpreted as unspecific fMRI signal drifts and were removed from all datasets by subtraction from the corresponding voxel-wise time series. The regressed ΔBOLD maps were averaged in consecutive non-overlapping 60 volume bins (corresponding to 1 min of acquisition) to reduce data dimensionality. Prior to analysis, spatial smoothing was applied using FSL with a Gaussian kernel (σ=2.5488, corresponding to a FWHM = 0.6 mm).

Quantification of ΔBOLD responses in the prefrontal cortex (i.e., Figure 1e) was performed using a linear mixed-effects model (*BOLD ∼ group × time + (1 + time | subject)*) implemented in R (lme4 package, with lmerTest extension) and fit using restricted maximum likelihood (REML). Fixed-effect statistics were obtained using t-tests with Satterthwaite’s approximation for degrees of freedom. Random intercepts captured between-subject differences in baseline signal, while random slopes modeled within-subject temporal dependencies across the time series.

Voxel-wise BOLD responses were quantified from mean ΔBOLD maps in three time windows: baseline (15 min), transition (15 min post-CNO), and CNO (20-35 mins post-CNO). Effects were assessed using a linear mixed-effects model (*BOLD ∼ group × time + (1 | subject)*) implemented in R (lme4), with random intercepts to account for between-subject variability.

#### fMRI connectivity

fMRI connectivity preprocessing was performed as described previously ^87^. The first 120 volumes of each time series were removed to account for gradient thermal equilibration artifacts. Data were then despiked, motion corrected, and co-registered to an in-house functional reference template. Motion traces and mean ventricle signal were regressed out. A band-pass filter was applied according to the dataset: 0.01-0.25 Hz for med-iso acquisitions and 0.01-0.1 Hz for halothane acquisitions. The resulting time series were spatially smoothed using AFNI (FWHM = 0.6 mm).

For fMRI functional connectivity analysis of ↑Excitation (both in sedated and awake animals) and ↓Inhibition experiments, we first qualitatively identified a time window of maximal efficacy. To this end, a moving correlation (movcorr function in MATLAB) was calculated between regions of interest (PFC and Retrosplenial) using overlapping windows of 900 volumes. The resulting time series were smoothed using a loess fit (30% overlap, MATLAB smooth function). Effect sizes between groups were then calculated (computeCohen_d, MATLAB), and voxel-wise FC differences were visually inspected to identify a time window of maximal effect (Figure S2).

Following the procedure described above, PFC voxel-wise connectivity was mapped within a window of maximal effect for each dataset (35-55 min for ↑Excitation, 15-25 min for ↑Excitation in awake mice, 15-35 min for ↓Inhibition, Figure S2). Voxel-wise FC changes were then quantified using a linear mixed-effects model implemented in R (lme4 package) in the form *FC(z) ∼ group × time + (1 | subject).* Time was treated as a categorical factor representing all phases (baseline, transition, and CNO windows). To account for potential baseline differences, subject-level variability was modeled using a random intercept. The resulting statistical interaction maps were cluster-corrected using FSL’s cluster function. Family-wise error rate (FWER) cluster correction was applied with a cluster-defining threshold of ∣t∣>2.1 and cluster significance threshold of p<0.05.

To plot the magnitude of FC changes between the PFC and retrosplenial cortex (e.g., Fig. 1e, 2e etc.), the mean fMRI signal across all voxels within each region of interest (Figure S6) was extracted, and Pearson correlations were computed within predefined post-CNO time windows (baseline, transition, 15-35 min, and 35–55 min). Correlations were Fisher z-transformed, and the same linear mixed-effects model employed in voxel-wise analyses was applied. P-values for each interaction term were corrected for multiple comparisons using the Benjamini-Hochberg procedure to control the false discovery rate (FDR, q=0.05).

To map brain-wide fMRI connectivity effects of PFC silencing, we fit a linear model to assess the effects of group, time, and their interaction on functional connectivity. Post hoc pairwise comparisons were performed using estimated marginal means (emmeans, R) to assess group differences within a post-CNO time window covering 15-30 min after CNO administration (to match the window used for multielectrode electrophysiological recordings in this dataset, see below). Voxel-wise statistical maps were FWER-corrected in FSL using a cluster-defining threshold of |t| > 2.1 and a cluster-level significance threshold of p<0.05. Functional connectivity between the PFC and retrosplenial cortex was statistically assessed within each time window using a two-sided t-test within each time window.

Voxel-wise quantifications of awake ↑Excitation experiment were carried out as described above. For each voxel, Fisher z-transformed functional connectivity (FCz) values were analyzed using a linear mixed-effects model of the form *FC (z) ∼ group × time + (1 | subject)*, fit voxel-wise. The group × time interaction term was then tested within the CNO window showing the strongest modulation (i.e. a 10 min window, starting 15 minutes after CNO injection, Figure S2). Voxel-wise statistical maps were FWER-corrected in FSL using a cluster-defining threshold of t<1.7 (reflecting the a priori hypothesis of decreased connectivity in DMN regions), and a cluster-level significance threshold of p<0.05. Region-of-interest quantification (PFC-thalamus) was performed by fitting the same model to Fisher z-transformed Pearson correlations computed from the two regional time series in the baseline and CNO windows.

To jointly evaluate the effects of ↑Excitation across awake and sedation datasets, a voxel-wise linear mixed-effects model was fit in R (lme4) of the form: *FC (z) ∼ group × time + anesthesia+ (1 | subject)*. Anesthesia was treated as a categorical fixed effect with two levels (anesthesia or awake), and within-subject variability was modeled with a random intercept. Time was defined as a categorical factor with two levels: baseline and CNO active window corresponding to maximal effect in each dataset (Figure S2). The resulting interaction maps were FWER-corrected in FSL using a cluster-defining threshold of |t|>2.1 and a cluster-level significance threshold of p<0.05. ROI quantifications were performed using the same model applied to Fisher-z values of the Pearson correlations in baseline and CNO active windows.

Quantification of the effect of SSp ↑Excitation and ↓Inhibition was performed using the same statistical model employed for the corresponding PFC manipulations. A linear mixed-effects model was fit voxel-wise across baseline, transition, and CNO active windows (the latter defined as a 15-min interval spanning 15-30 min post-injection). Interaction maps were FWER-corrected using the cluster function in FSL, with a cluster-defining threshold of |t|>2.1 and a cluster-level significance threshold of p<0.05. Functional connectivity between left and right SSp was quantified using the same model, with p-values corrected for multiple comparisons using the Benjamini– Hochberg procedure (q = 0.05).

Pharmacological fMRI response to PCP was mapped and quantified as previously described ^41^. Briefly, time series were spatially normalized to a common reference space and signal intensity changes were expressed as fractional BOLD. The first 10 volumes were discarded, and individual fMRI volumes were averaged into 36 s bins (30 volumes) to reduce data dimensionality. Voxel-wise group statistics were performed with a boxcar design in FEAT (v5.63) on spatially smoothed data (FWHM=0.6 mm). Voxel-wise group comparisons (PCP vs. vehicle) were performed in FSL using FLAME mixed-effects analysis with multilevel Bayesian inference. The resulting statistical maps were FWER-corrected in FSL using a cluster-defining threshold of |t| > 2.7 and a cluster-level significance threshold of p < 0.05. Voxel-wise functional connectivity differences between PCP and controls (vehicle) in the PFC were quantified using a two-tailed Student’s t-test followed by FWER cluster correction (as implemented in FSL) with a cluster defining threshold of |t|>2.1, and a cluster significance of p<0.05.

### Electrophysiological recordings

Electrophysiological ↑Excitation and ↓Inhibition studies were conducted under the same animal preparation and sedation regime adopted for fMRI acquisitions. Mice were anesthetized with isoflurane (4% induction, 2% maintenance), intubated, artificially ventilated, and head-fixed in a stereotaxic apparatus (Kopf). The tail vein was cannulated for subsequent medetomidine infusion, and an additional intraperitoneal cannula was placed for CNO administration. Two independent single-shank, 16-channel linear electrodes (Neuronexus, USA; A1×16-1.5mm-50-177) were inserted simultaneously into the PFC (from Bregma: AP +1.7 mm, ML ±0.3 mm, DV-1.7 mm) and retrosplenial cortex (from Bregma: AP -2.5 mm, ML ±0.3 mm, DV -0.8 mm) at a rate of 1 µm/min, matching the stereotaxic coordinates used for viral injection. The distance between the two probes was ∼2.5 mm. A silver wire was placed over the cerebellum for reference. Following electrode insertion, a bolus of medetomidine (0.05 mg/kg) was administered intravenously, followed 5 min later by a continuous infusion at a rate of 0.1 mg/kg. Isoflurane concentration was then reduced to 0.5% to maintain a combined medetomidine-isoflurane sedation regimen consistent with that used in fMRI studies. Recordings commenced ≥45 min after isoflurane cessation to allow complete washout of induction anesthesia.

The experimental protocol for the Silencing experiment is described in detail elsewhere ^16^. Briefly, animals were anesthetized with isoflurane (4% induction, 2% maintenance), intubated, and head-fixed in a stereotaxic apparatus. The tail vein was cannulated for CNO administration. Surgeries and electrode insertion were performed as described above. Following electrode insertion, isoflurane was discontinued and replaced with halothane (0.8%). Electrophysiological recordings commenced ≥45 min after the halothane switch to ensure complete washout of isoflurane.

Electrophysiological signal was next recorded to cover a 15 min pre-injection time window, and a 60 min post CNO administration. Extracellular voltage signals were amplified using an RHD2000 system (Intan Technologies; RHD Recording Controller Software v2.09) and sampled at 20 kHz.

### LFP and multi-unit activity (MUA)

To extract local field potentials (LFP), the raw extracellular signal was first down-sampled to 4 kHz and band-pass filtered using a two-step procedure ^88^. A 4th-order low-pass Butterworth filter with a cut-off frequency of 1 kHz was initially applied to the 4 kHz signal. The resulting signal was then down-sampled to 2 kHz and filtered with a Kaiser window filter (passband 0.1–250 Hz, sharp transition bandwidth 1 Hz, passband ripple 0.01 dB, stopband attenuation 60 dB). Finally, the signal was down-sampled to 1 kHz. All filters were applied forward and backward to prevent phase distortions.

Multi-unit activity (MUA) was computed as described previously^88^. The extracellular signal was first high-pass filtered (4th order Butterworth filter with cut off frequency over 100Hz), then band-pass filtered between 400-3000 Hz using a Kaiser window filter (transition bandwidth 50 Hz, stopband attenuation 60 dB, passband ripple 0.01 dB). Spikes were detected from the high-frequency signal using a thresholding method^89^, with the threshold set at four times the median of the absolute signal, divided by 0.6745, calculated over the entire time series (baseline and post-CNO combined). Events exceeding this threshold were considered candidate neuronal spikes. Only events separated by ≥1 ms were retained.

To assess the effect of chemogenetic manipulations, spiking activity was computed in experimental and control animals as described above and segmented into 1-minute bins. We verified that this bin size yielded largely temporally independent firing-rate samples, with negligible temporal dependence between one sample and the next. This was verified by computing the autocorrelation function of the binned firing rate and by confirming that the lag at which autocorrelation dropped below the 95^th^ percentile of the distribution expected under the no-autocorrelation hypothesis (obtained from the *bounds* output of the MATLAB function *autocorr*) was consistently <1 min^16^. For each 1-minute bin, we computed the channel-averaged firing rate and standardized it by subtracting the mean firing rate during the baseline (pre-CNO) period and dividing by the corresponding standard deviation. Abrupt changes in standardized firing rate (>8× the mean of the preceding 10 time points) were classified as non-biological artifacts and removed; the resulting gaps were linearly interpolated in MATLAB. Standardized firing rates were pooled across subjects and analyzed in two post-CNO windows (15-35 min and 35-55 min). Group differences in median firing rate were assessed using two-sided Wilcoxon rank-sum tests, with p-values FDR corrected for multiple comparisons using the Benjamini-Hochberg procedure (q = 0.05). For all subsequent analyses (LFP power and coherence), our investigations were restricted to the post-CNO time window previously identified as the period of maximal effect in the corresponding fMRI dataset, unless otherwise stated.

### Spectral analysis

LFP power spectra were computed using the *pspectrum* function in Matlab with frequency resolution equal to 1Hz. To quantify the effect of CNO on LFP rhythms, we computed a power spectrum modulation index as follows. For each subject, we first computed the channel-averaged power spectrum during the baseline recording. We then computed channel-averaged power spectra in consecutive 1-min bins within the post-CNO window. As for the firing rate analyses, the bin duration was informed by the autocorrelation function of the spectrogram: we verified that the autocorrelation fell within the 95^th^ percentile of the distribution expected under the no-autocorrelation hypothesis (see above), indicating that the corresponding 1-min spectral estimates exhibited negligible temporal dependence. Then, for each one-minute bin after injection, we defined the modulation index as the relative difference between channel-averaged power spectrum after injection and baseline (i.e., their difference divided by their sum),

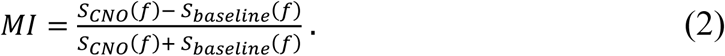

This modulation index ranges between -1 and 1 and describes the changes due to drug injection for each frequency.

To statistically assess CNO-related group differences in the power spectrum while controlling for multiple comparisons across frequencies, we used a frequency-resolved cluster-based correction approach^90^. At each frequency, spectra from control and experimental animals were compared using a two-sample t-test. The resulting t-statistics were thresholded at the critical value corresponding to α = 0.05 (Student’s t inverse cumulative distribution), and contiguous suprathreshold frequency bins were grouped into clusters. For each cluster, we computed a cluster-mass statistic defined as the sum of t-values within the cluster.

To determine the statistical significance of frequency clusters we used a permutation test ^90^. We generated 10^4^ permutations shuffling group labels of the modulation index values, and we repeated the statistical test described above; the largest cluster mass value for each permutation was used to define the null distribution of cluster mass values. Then, a p-value for each cluster obtained in the original data was defined as the ratio between the number of mass values in the null distribution larger than the cluster mass values, divided by the number of permutations.

### Multielectrode LFP coherence

We computed the magnitude-squared coherence between channel-averaged LFP signals of two areas using the *mscohere* Matlab function (that uses Welch’s overlapped averaged periodogram method) with 2000-sample windows and 1000-sample overlap^91^. Coherence was estimated in consecutive 1-min bins; for each bin, we computed a coherence modulation index defined as in Eq. (2), (*C*_*CNO*_ − *C*_*baseline*_) /*C*_*CNO*_ + *C*_*baseline*_ where *C*_*CNO*_ is coherence after injection and *C*_*baseline*_ is baseline coherence.

Coherence modulation was evaluated only within frequency ranges in which baseline coherence was statistically significant. Significance thresholds were determined using a bootstrap approach: coherence was computed for 10,000 shuffled time series to generate a null distribution at each frequency, and the significance threshold was defined as the mean of the 95th percentile of the null distribution across frequencies. This procedure identified significant coherence for frequencies <40 Hz in all animals and groups; faster frequencies were not significant and were excluded from further analyses. To obtain a statistical assessment of CNO effects on coherence across groups, we carried out a frequency-resolved analysis, using the same procedures described above for the power spectrum. Finally, to summarize effects within frequency ranges showing significant coherence (slow through β bands), we computed for each subject in each 1-min bin a band-wise modulation index by averaging the frequency-resolved modulation indices within the band limits (slow 0.1-1 Hz, δ 1-4 Hz, θ 4-8 Hz, α 8-12 Hz, and β 12-30 Hz). Group differences in these band-wise indices were assessed using two-sided Wilcoxon rank-sum tests, with p-values corrected for multiple comparisons using the Benjamini-Hochberg procedure (q = 0.05).

To assess interareal coherent changes in ultra-slow fluctuations of the γ-band envelope of the LFP, we followed the procedure described in^19^. LFP spectrograms were computed for the last 10 min of baseline and the post-CNO window, using the same procedure described above. The γ-band envelope was calculated as the integrated spectrogram power over 30-70 Hz. Coherence was then computed between the γ-band envelopes extracted from the two regions. Chemogenetic effects were quantified using the modulation index described above. Between-group comparisons were performed using two-sided Wilcoxon rank-sum tests, with p-values corrected for multiple comparisons using the Benjamini-Hochberg procedure (q = 0.05).

Plots were generated using Prism Graphpad 9.1. Plot truncation at extremities is introduced by the software to avoid representation of kernel density-related fictitious values extending beyond the observed range.

### Neural network simulations

We performed several simulations of different neural networks. All simulations have a basic single-area circuit module which is either simulated on its own (i.e., Figure S1) or combined in a model of three interacting areas (i.e., Figure 6).

#### Single area module (for Figure S1)

The basic single-area circuit module was composed of two recurrently connected excitatory-inhibitory (E-I) populations. Each single-area module was implemented as a recurrent model of leaky integrate-and-fire excitatory pyramidal (E) and inhibitory (I) neurons, with architecture and parameters taken from prior work ^30, 36, 92, 93^ (Table S1). Each circuit comprised 5,000 neurons, of which 4,000 were excitatory pyramidal cells forming AMPA-like synapses, and 1,000 were inhibitory interneurons forming GABA-like synapses. Within each circuit, neurons had identical intrinsic properties and were connected with an intra-circuit random connection probability of 0.2 ^33, 92, 93^(see Table S2). When simulated with a stationary input, these models are known to intrinsically generate only gamma-band oscillations due to interactions between E and I neurons^34, 94^. When instead simulated with inputs fluctuating at low frequencies, these networks maintain the internal generated gamma oscillations, and in addition exhibit low-frequency oscillatory activity reflecting entrainment to the inputs^34^. For the study of the single-area module in Supplementary Figure 1 we started form a reference set of parameters (reported in Tables S1-2). In Figure S1B we varied the resting potential of E neurons as indicated on the x axis. In Figure S1C we varied the conductance of excitatory to excitatory recurrent connections *g_EE_*as indicated on the x axis.

#### Three area model (i.e., Figure 6)

The three-area model was constructed by connecting three single-area modules as follows. We connected two "sending” single area modules via direct synaptic connections from the E neurons of the sending area to the E neurons of the third “receiving” module. The interareal connection probability was set lower than the intra-area probability (0.05 vs. 0.2), reflecting the relative sparsity of long-range cortico-cortical connections compared with local recurrent circuitry^95, 96, 97^. The synaptic conductance of inter-are E connections had the same value as those of intra-area E connections. The receiving area represents RS cortex, which is the area for which we measure the effect in PFC-RS coherence changes when PFC E-I balance is manipulated. The first sending area models the PFC and is the one whose E-I parameters are manipulated to mimic the chemogenetic manipulation. The second area sending to RS is added to model the fact that RS has inputs from many areas, some of which will not be affected by the PFC manipulation.

All three single-area modules received an external input, representing influences from other brain regions. Each source of external input was modeled as a set of input spikes generated by 4,000 Poisson input neurons that projected via AMPA-like synapses to both the excitatory and inhibitory populations in each circuit with a connection probability of 0.2. This set of input spikes has two components. The first component consisted of stationary Poisson firing with a given mean firing rate, which determines the average depolarization of the neurons within each module. This component was generated independently for each of the three circuits and models area-specific inputs. The second had a non-stationary Poisson firing with a slowly varying firing rate set by an Ornstein-Uhlenbeck (OU) process with a time constant of 80 ms (which thus generates most power at frequencies slower than 4 Hz), capturing slow fluctuations correlated over wide ranges typical of spontaneous cortical state changes and thus shared by several cortical areas. These connected networks transmit information through firing rates in the directly connected areas, and oscillatory influences of these direct information transmission are confined to high-frequency coherence ^47^.

To model the in *vivo* effect of PFC chemogenetic manipulations, we ran three sets of simulations, with parameters adjusted independently to reproduce the corresponding experimental conditions. All manipulations were restricted to the PFC module leaving the RS receiving area and the second sending area unaltered. In the first set, the hSYN-hM4Di condition was mimicked by hyperpolarizing both excitatory and inhibitory PFC neurons (resting membrane potential from-70 to-73 mV) and proportionally reducing synaptic strengths by 25%. In the second set, the excitatory DREADD CamkII-hM3D(Gq) manipulation was simulated by depolarizing excitatory neurons in PFC (resting membrane potential −70 mV to −69 mV). In the third set, the inhibitory PV-hM4Di manipulation was implemented by hyperpolarizing inhibitory neurons in PFC (resting membrane potential −70 mV to −71 mV) and proportionally reducing inhibitory synaptic strength (g_IE_ and g_II_) by 10%. In each experiment, the control condition was defined by identical neuronal and synaptic default parameters across all three experiments. In contrast, the parameters of the stationary input differed across control conditions. In the CaMKII-hM3D(Gq) and PV experiments, they were adjusted to reproduce the gamma oscillations observed in the experimental control data by increasing the mean of the depolarizing OU input to 3.0 spikes/s per neuron.

All simulations lasted 60.5 seconds, with the first 0.5 s discarded to allow the network to reach a stationary regime. Regional LFPs were computed as the sum of the absolute AMPA- and GABA-mediated postsynaptic currents onto excitatory neurons, consistent with previously validated LFP proxies^34, 48, 98^. LFP signals were sampled at 1 kHz and segmented into non-overlapping time-bins, which were used as trial units for coherence and firing-rate analyses. Similar to the analysis of real LFP data, magnitude-squared coherence between LFP time series was computed for each trial using Welch’s method with Hamming windows. Window length was set adaptively to include at least three cycles of the center frequency, with 50% overlap. Coherence was then averaged within the same canonical frequency bands used for the in vivo analyses. Differences between control and manipulation conditions were assessed using two-sided Wilcoxon rank-sum tests, with p-values corrected for multiple comparisons using the Benjamini–Hochberg procedure (q = 0.05).

Population firing rates were computed from simulated spike activity for both control and manipulation conditions, using the same non-overlapping 20-s trial segmentation adopted for coherence analyses. Fore each time-bin, the average population firing rates were computed separately for E and I neurons, and averaging over all neurons. As in empirical electrophysiological recordings, to facilitate comparisons across conditions, firing rates were z-scored relative to the control distribution (using the control mean and standard deviation), and group differences were tested using two-sided Wilcoxon rank-sum tests. The resulting p-values were adjusted for multiple comparisons using the Benjamini-Hochberg false discovery rate correction (q = 0.05).

### Electrophysiological correlates of fMRI connectivity changes

To maximize comparability across datasets for multimodal correlation analyses, electrophysiological and fMRI metrics were extracted from matched post-CNO windows selected to reduce sensitivity to non-specific drug effects and anesthesia-related drifts, while providing sufficiently long segments for stable estimation. We thus used a 20 min window starting 20 min after CNO injection for both ↑Excitation and ↓Inhibition datasets (a window central to the peak effects observed in those manipulations), and a 15-30 min post-CNO window for the silencing dataset. The latter choice was constrained by the duration of our prior multielectrode electrophysiological recordings, which for some subjects did not consistently extend beyond 30 min post-injection. We next jointly analyzed the electrophysiological correlates of all manipulations (↑Excitation, ↓Inhibition, Silencing, and their respective controls), by computing the Spearman correlation between PFC band-limited power modulation and fMRI connectivity modulation, defined as:

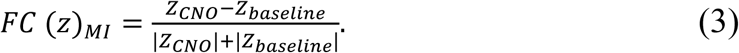

where *Z* denotes Fisher z–transformed functional connectivity. Correlations were computed between PFC-Retrosplenial cortices in the ↑Excitation and ↓Inhibition experiments, and between PFC-Retrosplenial cortices as well as PFC-Thalamus in the Silencing experiment, for which three-electrode recordings were available. In these analyses, each data point corresponds to the group-averaged LFP modulation within a given frequency band (slow 0.1-1 Hz, δ 1-4 Hz, θ 4-8 Hz, α 8-12 Hz, β 12-30 Hz, γ 30-70 Hz, high γ 70-150 Hz) or MUA, and the group-averaged fMRI connectivity modulation between the corresponding regions of interest. P-values were corrected for multiple comparisons using the Benjamini–Hochberg procedure (q = 0.05). Similarly, for power coherence modulation, we calculated the Spearman correlation between LFP coherence modulation across electrode pairs (PFC-Retrosplenial in ↑Excitation and ↓Inhibition experiments, and PFC-Retrosplenial and PFC-Thalamus in the Silencing experiment) and the corresponding fMRI connectivity modulation across the same region pairs. We restricted statistical testing to frequency bands in which LFP coherence and connectivity modulation exhibited concordant signs (slow, δ and 0.1-4 Hz), as these bands could provide a biologically plausible explanation for the observed effects. We reported untested Spearman values (Figure 5) only for reference. P-values were corrected for multiple comparisons using Benjamini-Hochberg FDR procedure (q = 0.05).

## Data availability

fMRI and electrophysiology data generated for this study will be available for download upon publication of the study.

## Code availability

The code used for preprocessing and analyzing mouse rsfMRI data is available at https://github.com/functional-neuroimaging/rsfMRI-preprocessing. Code for analyses of electrophysiological recordings is available at https://github.com/panzerilab/Ephys-Analyses_pub. Code for the Simulations is available at https://github.com/panzerilab/Electrophysiologically-defined-excitation-inhibition-autism-neurosubtypes

## Acknowledgments

This work has been funded by the European Research Council (ERC) under the European Union’s Horizon 2020 research and innovation program (no. 101125054 #BRAINAMICS and no. 802371 #DISCONN to A.G.), the Simons Foundation (SFARI; grant number 982347 to A.G and S.P.), the Brain and Behavior Research Foundation (NARSAD; Independent Investigator Grant; no. 25861, to A.Go.), the EU Marie Sklodowska-Curie Action (101152984 to S.B.M.), and by the Brain and Machines Flagship Program of the Italian Institute of Technology.

