## Supplementary Figures for "Cortical excitability inversely modulates fMRI connectivity via low-frequency neuronal coupling"

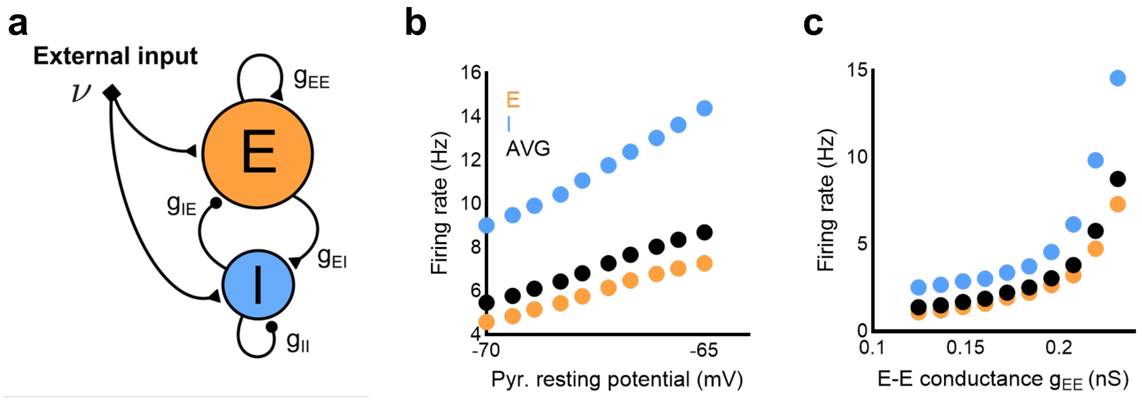


**Supplementary Figure 1. Firing rate of E and I neurons changes in a coordinated way in recurrently coupled E-I networks. (a)** Schematic of the simulated individual-area recurrent E-I network. The model consists of two neuronal populations: 4000 excitatory (E) pyramidal neurons and 1000 inhibitory (I) interneurons. Neurons are connected through randomly structured intra- and inter-population synaptic connections. Both populations receive an external input which determines the baseline excitability of individual neurons. **(b)** Changes in the firing rates of E (orange), I (blue) neurons and their average (black) when the pyramidal neuron excitability is manipulated by changing the resting membrane potential of excitatory neurons. Each point represents the mean firing rate averaged across two independent 4-s simulation runs. **(c)** Same analysis as in **(b)** but plotting changes in firing rate as function of the strength of the synaptic efficacy of recurrent excitatory connections ($g_{EE}$). In both simulation sets, excitatory and inhibitory firing rates change in a coordinated manner; for example, increasing the intrinsic excitability of excitatory neurons also increases inhibitory-neuron firing through recurrent coupling. Thus, the mean network firing rate can be used as a practical index of the operating point of the E-I circuit.

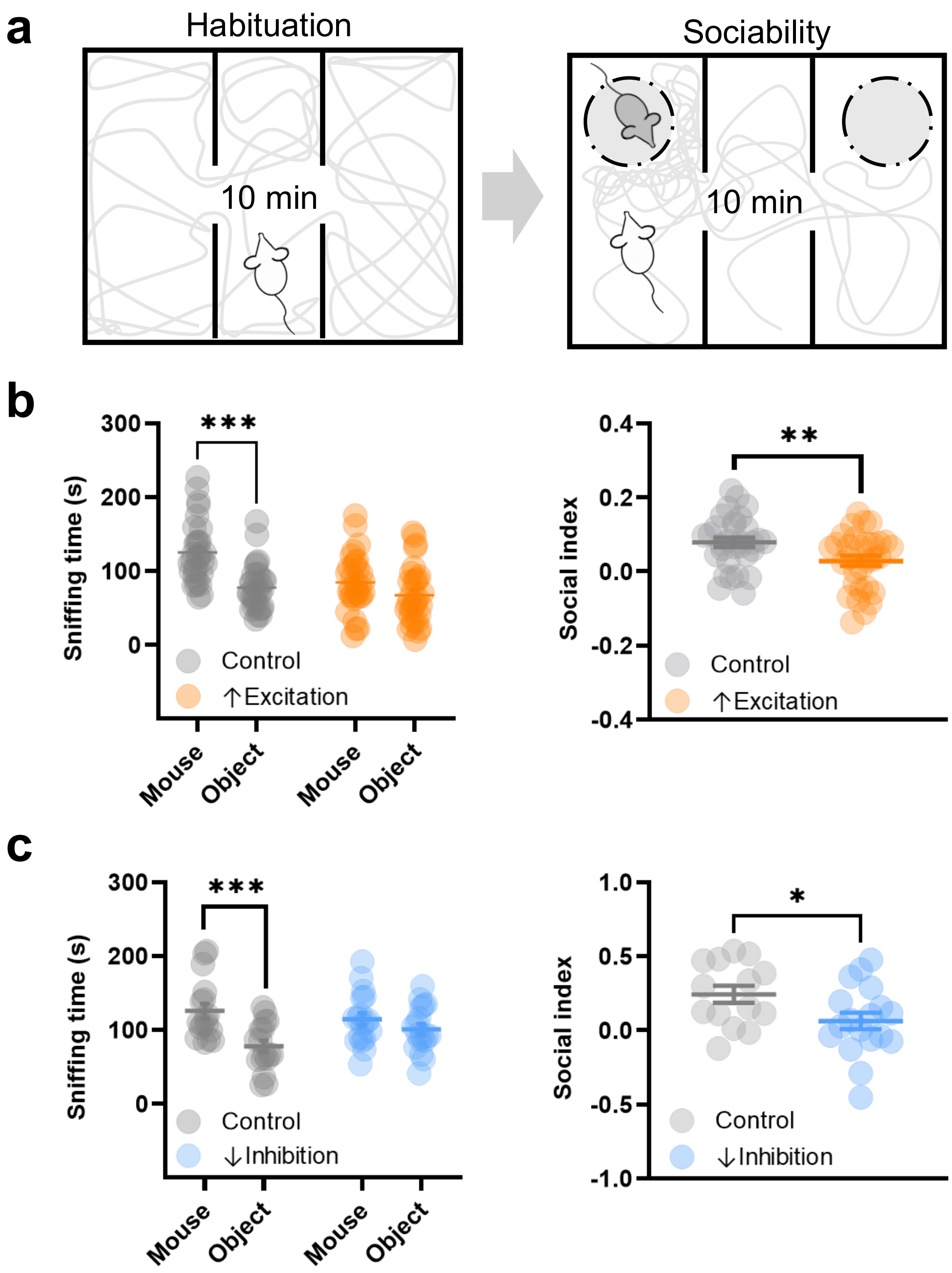


**Supplementary Figure 2. Increasing PFC excitability alters sociability in the three-chamber test. (a)** Three chamber sociability paradigm (habituation and sociability phases, 10 min each). **(b)** Sniffing time for the social stimulus (mouse) versus object, and corresponding sociability index for ↑Excitation (n = 31) and control (n = 31) animals. **(c)** Sniffing time for the social stimulus (mouse) versus object, and corresponding sociability index for ↓Inhibition (n = 18) and control (n = 14) mice. Data are shown as mean ± SEM; *p<0.05, **p<0.01, ***p<0.001, Student's t test.

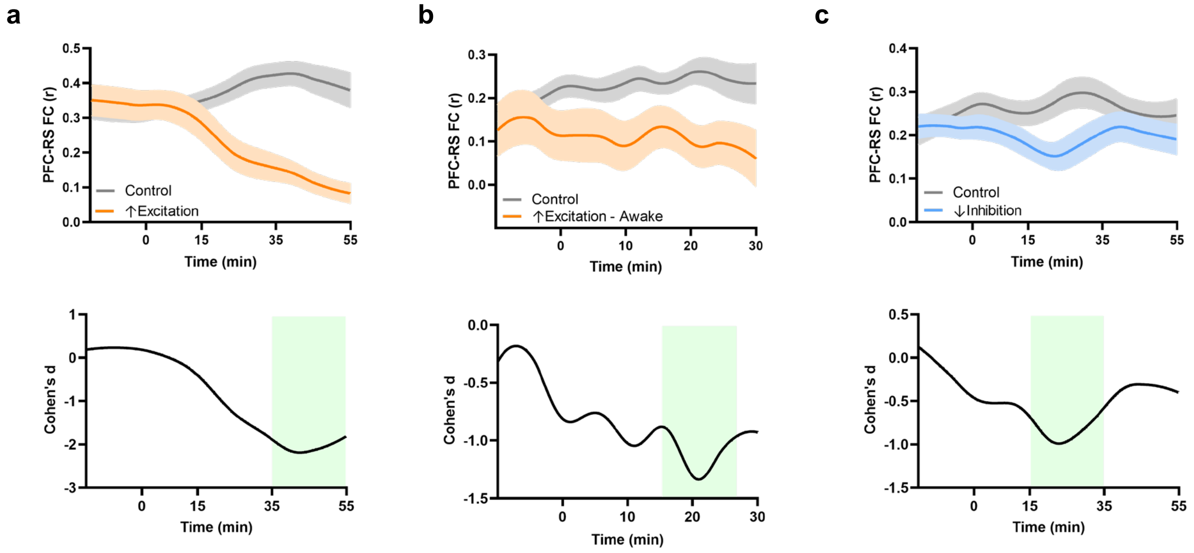


**Supplementary Figure 3. Temporal evolution of PFC–RS connectivity in ↑Excitation and ↓Inhibition experiments for the newly acquired datasets.** Moving correlation of PFC and RS fMRI connectivity (FC, top; mean ± SEM) and corresponding effect size (bottom; Cohen's d) for **(a)** ↑Excitation (med-iso sedation), **(b)** ↑Excitation (awake) and **(c)** ↓Inhibition (med-iso sedation). Green rectangles mark time-windows of maximal effect used to map fMRI connectivity differences (35-55, 15-25 and 15-35 min post CNO, for ↑Excitation, ↑Excitation awake, and ↓Inhibition, respectively). PFC: prefrontal cortex; RS: retrosplenial cortex.

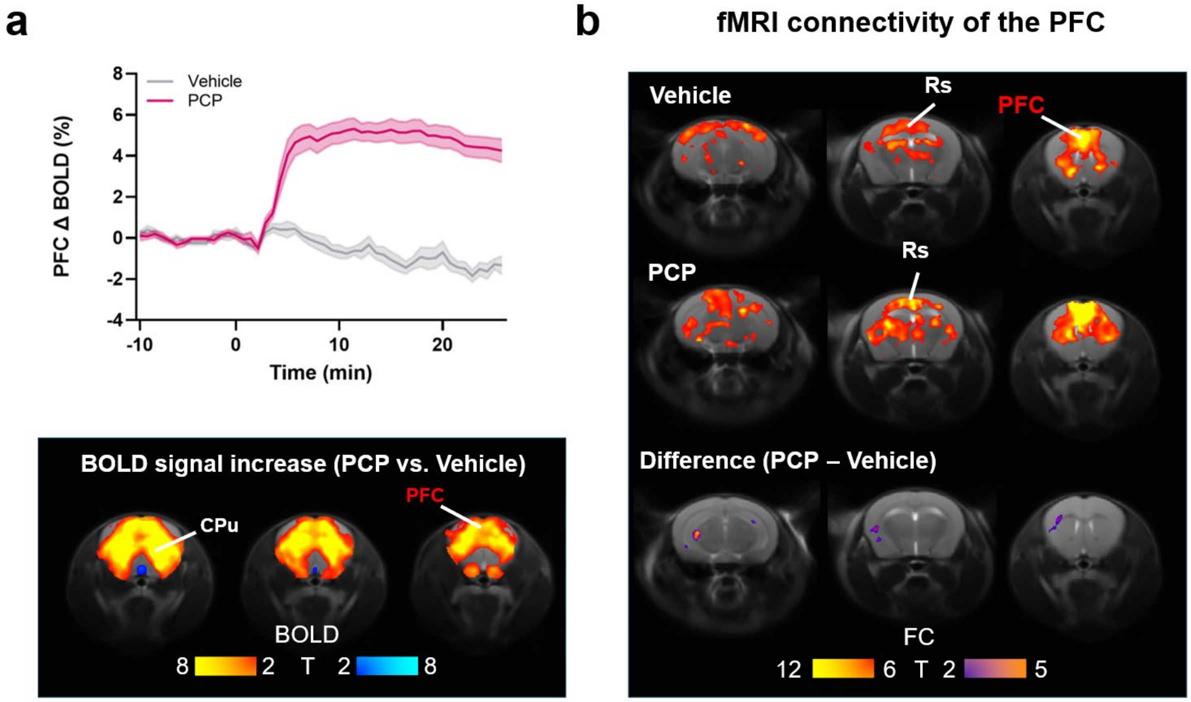


Supplementary Figure 4. PCP-evoked BOLD response and functional connectivity in the PFC. **(a)** Mean ΔBOLD time course in the PFC after vehicle or PCP administration (1 mg/kg; n = 10 per group; mean ± SEM) and corresponding voxel-wise group contrast (PCP-vehicle) mapped using GLM and a boxcar regressor ( $|t|>2.7$; FWER-corrected, cluster $p<0.05$). **(b)** Seed functional connectivity maps for the PCP in vehicle and PCP-treated groups, and corresponding voxel-wise difference map ($|t|>2.1$; FWER corrected; $p<0.05$). CPu, caudate-putamen; FC, functional connectivity; PCP, phencyclidine; PFC, prefrontal cortex; RS, retrosplenial cortex.

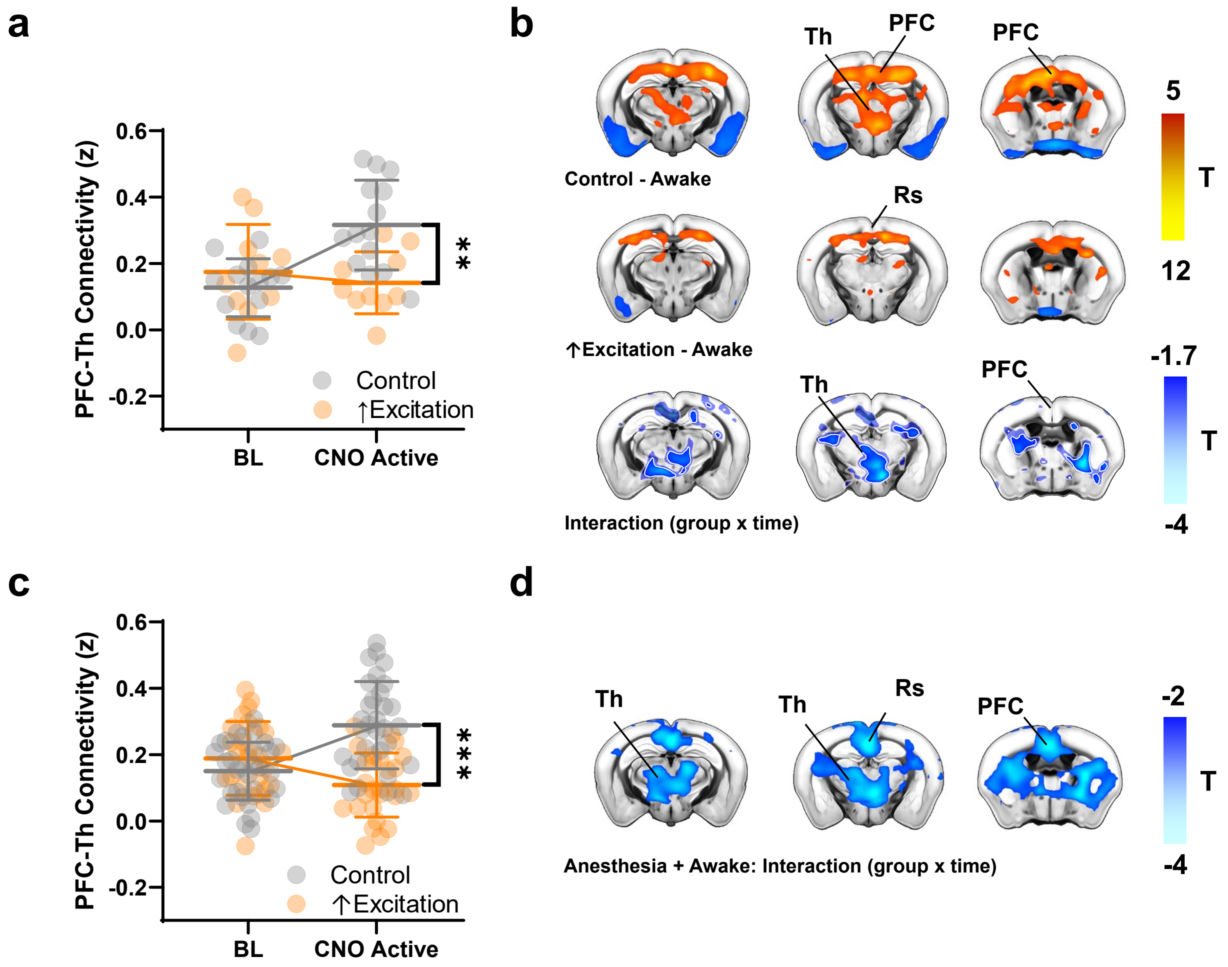


**Supplementary Figure 5. fMRI connectivity effects in awake mice, and combined anesthesia + awake analysis.** **(a)** PFC–thalamus connectivity in ↑Excitation (n = 10) versus control mice (n = 14) acquired under awake conditions (BL, baseline pre-CNO; CNO active, 15-25 min post-CNO; group × time interaction; linear mixed-effects model; **p<0.01). **(b)** PFC-seed-based fMRI connectivity map in ↑Excitation and control animals, and corresponding reduced connectivity voxelwise maps (group x time interaction, linear mixed-effects mode, cluster-FWER corrected t<1.7; cluster p<0.05). **(c)** PFC-thalamus fMRI connectivity in the combined awake + sedation ↑Excitation analysis, quantified in window of maximal activity (CNO active; ↑Excitation n = 27; controls n = 31; group × time; linear mixed-effects model; ***p<0.001). **(d)** Corresponding voxel-wise interaction map (|t|>2.1; cluster-FWER corrected; cluster p<0.05). Data in **b** shows FWER-corrected clusters (white edge) overlaid on the non-corrected maps (transparent blobs). BL: baseline, PFC: prefrontal cortex, Rs: retrosplenial cortex, Th: thalamus.

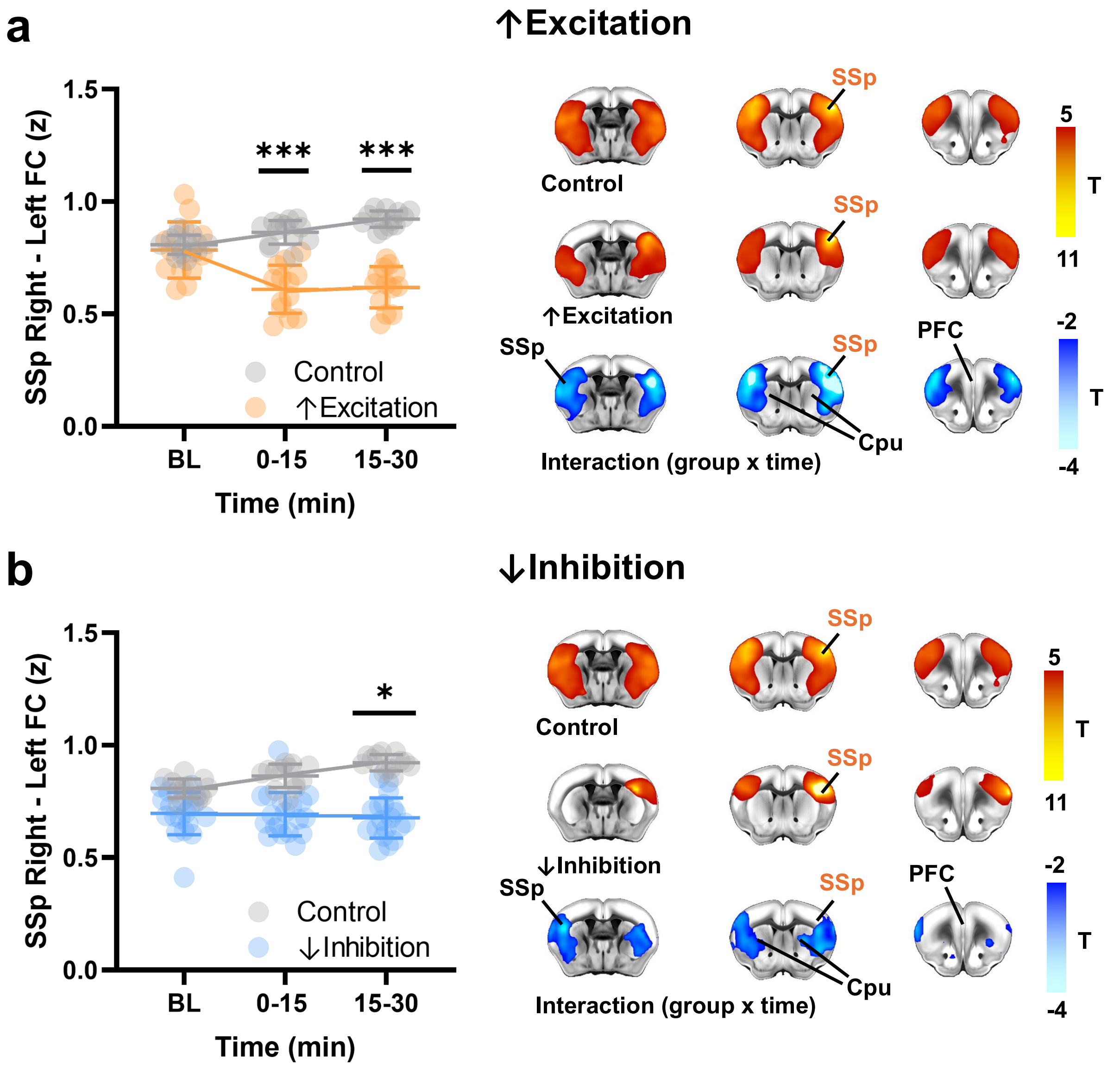


**Supplementary Figure 6. Increasing excitability of somatosensory cortex results in reduced fMRI connectivity. (a)** Quantification of homotopic bilateral SSp connectivity (left) and SSp connectivity maps (right) for SSp ↑Excitation (n = 13) versus controls (n = 13), mapped in the 15-30 min post-CNO window. **(b)** Quantification of homotopic bilateral SSp connectivity (left) and SSp seed connectivity maps (right) for ↓Inhibition (n = 19) versus controls (n = 13), mapped in the 15-30 min post-CNO window. Difference maps in **(a)** and **(b)** show the group × time interaction from a linear mixed-effects model, cluster-FWER corrected ($|t|>2.1$; cluster $p<0.05$). Line plots show mean ± SD. *$p<0.05$; ***$p<0.001$ for the interaction term (linear mixed-effects model; FDR-corrected). SSp, primary somatosensory cortex; CPu, caudate-putamen; FC, functional connectivity; BL, baseline.

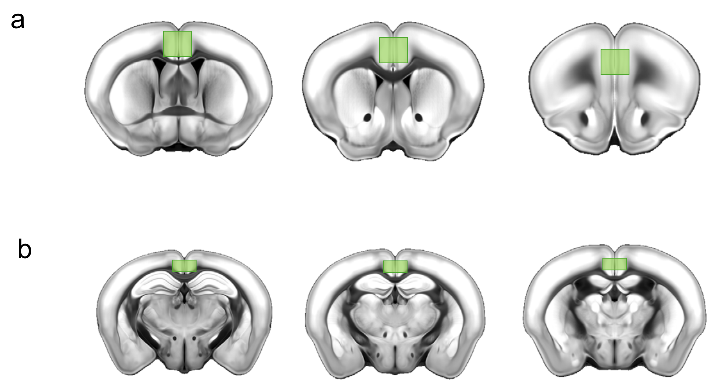


**Supplementary Figure 7**. Anatomical location of multi slice ROIs used for regional quantification of effects. Coronal sections illustrating the PFC ROI (a) and RS ROI (b) used for ROI-based connectivity and regional effect quantification.
