## Supplementary Table I for "Cortical excitability inversely modulates fMRI connectivity via low-frequency neuronal coupling"

| A: Model summary | |
|---|---|
| Structure | Excitatory-inhibitory (E-I) network |
| Populations | One excitatory and one inhibitory |
| Input | 2 independent external input populations: one delivering Poisson spike trains and one delivering spike trains with rates governed by an Ornstein–Uhlenbeck (OU) process. |
| Measurement | Spikes, AMPA and GABA currents for the two excitatory populations |
| Neuron model | Cortex: leaky integrate-and-fire (LIF) with fixed threshold and fixed absolute refractory time |
| Topology | None |
| Connectivity | Random and sparse |

| B: Populations | | |
|---|---|---|
| Type | Elements | Size |
| Pyramidal cells | LIF neurons | 4000 |
| Interneurons | LIF neurons | 1000 |
| Thalamic Input | Homogeneous Poisson generator | 1 |
| Slow-cortical input | Inhomogeneous Poisson generator | 1 |

| C: Connectivity Intra-area | | | |
|---|---|---|---|
| Name | Source | Target | Pattern |
| Recurrent E-E | Pyramidal | Pyramidal | Random convergent (p=0.2), weight = $g_{EE}$ |
| Recurrent E-I | Pyramidal | Interneurons | Random convergent (p=0.2), weight = $g_{EI}$ |
| Recurrent I-E | Interneurons | Pyramidal | Random convergent (p=0.2), weight = $g_{IE}$ |
| Recurrent I-I | Interneurons | Interneurons | Random convergent (p=0.2), weight = $g_{II}$ |
| External Ex-E | External pop. 1 | Pyramidal | Random convergent (p=0.2), weight = $g_{EXE}$ |
| External Ex-I | External pop. 1 | Interneurons | Random convergent (p=0.2), weight = $g_{EXI}$ |
| External SF-E | External pop. 2 | Pyramidal | Random convergent (p=0.2), weight = $g_{SFE}$ |
| External SF-I | External pop. 2 | Interneurons | Random convergent (p=0.2), weight = $g_{SFI}$ |

| E: Neuron model | |
|---|---|
| Type | Leaky integrate-and-fire |
| Description | $\tau_m \frac{dV(t)}{dt} = -V(t) + V_{leak} - \frac{I_{tot}(t)}{g_{leak}}$,<br><br>$I_{tot}(t) = \sum_{N_{EXC_{rec}}} I_{EXC_{rec}}(t) + \sum_{N_{INH_{rec}}} I_{INH_{rec}}(t) + I_{EXC_{ext}}(t)$. |

| F: Synapse model | |
|---|---|
| Type | Conductance-based synapse, difference of exponentials |
| Description | $I_{syn}(t) = g_{syn} S_{syn}(t)(V(t) - E_{syn})$,<br>If a presynaptic spike occurs:<br>$s_{syn}(t) = \frac{\tau_m}{\tau_d - \tau_r}\left[exp\left(\frac{-t-\tau_l}{\tau_d}\right) - exp\left(\frac{-t-\tau_l}{\tau_r}\right)\right]$ |

| G: Input | | |
|---|---|---|
| Name | Type | Description |
| Depolarization input | Poisson spikes | Stationary spike trains, mean $\nu_T$ |
| Slow fluctuating input | Poisson spikes | Non-stationary spike trains, Poisson mean $\nu_c$ modulated by OU process |

| H: Global Simulation parameters | |
|---|---|
| Simulation duration | 4.5 s |
| Temporal resolution | 0.1 $ms$ |
| Startup transient | 500 $ms$ |

**Supplementary Table 1: Description of the architecture and simulation settings of the single area neural network module.** Overview of the single-area network module architecture and simulation settings, encompassing population structure, connectivity, neuron and synapse models, and global simulation parameters used in the *in-vitro* simulations of the single-module network. This is the basic module that is used in Supplemental Figure 1 on its own, and that is used for each of the 3 single areas modules that are made to interact in Figure 6 (with inter-area interaction parameters described in Methods).
