## Supplementary Table II for "Cortical excitability inversely modulates fMRI connectivity via low-frequency neuronal coupling"

| A: Neuron parameters | | | | |
|---|---|---|---|---|
| Parameter Name | Symbol | Pyramidal cells | Interneurons | Unity of measure |
| Leak (resting) membrane potential | $V_{leak}$ | -70 | -70 | $mV$ |
| Spike threshold potential | $V_{threshold}$ | -52 | -52 | $mV$ |
| Reset potential after a spike | $V_{reset}$ | -59 | -59 | $mV$ |
| Membrane refractory period | $\tau_{refractory}$ | 2 | 1 | $ms$ |
| Leak conductance | $g_{leak}$ | 25 | 20 | $nS$ |
| Membrane capacitance | $C_m$ | 500 | 200 | $pF$ |
| Membrane time constant | $\tau_m$ | 20 | 10 | $ms$ |
| **B: Connection parameters** | | | | |
| Parameter Name | Symbol | Pyramidal cells | Interneurons | Unity of measure |
| AMPA synaptic reversal potential | $E_{AMPA}$ | 0 | 0 | $mV$ |
| GABA synaptic reversal potential | $E_{GABA}$ | -80 | -80 | $mV$ |
| AMPA synapse rise time constant | $\tau_{r(AMPA)}$ | 0.4 | 0.2 | $ms$ |
| AMPA synapse decay time constant | $\tau_{d(AMPA)}$ | 2 | 1 | $ms$ |
| GABA synapse rise time constant | $\tau_{r(GABA)}$ | 0.25 | 0.25 | $ms$ |
| GABA synapse decay time constant | $\tau_{d(GABA)}$ | 5 | 5 | $ms$ |
| Latency of intra-area transmission | $\tau_{l(intra)}$ | 1 | 1 | $ms$ |
| Latency of feed-forward connections | $\tau_{l(FF)}$ | 2 | 2 | $ms$ |
| Recurrent AMPA synaptic conductance | $g_{AMPA(rec.)}$ | 0.178 | 0.233 | $nS$ |
| External depolarizing AMPA synaptic conductance | $g_{AMPA(ext.)}$ | 0.234 | 0.317 | $nS$ |
| Slow fluct. input AMPA synaptic conductance | $g_{AMPA(slow.)}$ | 0.187 | 0.254 | $nS$ |
| GABAergic synaptic conductance | $g_{GABA}$ | 2.01 | 2.7 | $nS$ |

| C: External input parameters | | | |
|---|---|---|---|
| External input type | Mean | Temporal Variability | Time constant |
| Depolarization input | 2 spikes/s/cell | - | — |
| Slow fluctuating input | 0 spikes/s/cell for Suppl Fig S1<br>0.3 spikes/s/cell for Fig 6 | 0.16 spikes/s/cell for fig S1<br>0.3 spikes/s/cell for Fig 6 | 16 ms for simulations in Suppl Fig 1 and 80 ms for simulations in Fig 6 |

**Supplementary Table 2: Biophysical parameters of the single area neural network module.** Summary of the neuronal and synaptic biophysical parameters, including membrane properties, synaptic kinetics, conductances, and external input statistics for the single area module used on its own in Suppl Fig 1 and used for each of the 3 single areas modules that are made to interact in Figure 6. For Figure 6, the parameters in the above table are those used for the control condition simulation. The parameters that were changed from this configuration to mimic the DREADD application are reported in Methods section.
